# Extracellular DNA promotes efficient extracellular electron transfer by pyocyanin in Pseudomonas aeruginosa biofilms

**DOI:** 10.1101/2019.12.12.872085

**Authors:** Scott H. Saunders, Edmund C.M. Tse, Matthew D. Yates, Fernanda Jiménez Otero, Scott A. Trammell, Eric D.A. Stemp, Jacqueline K. Barton, Leonard M. Tender, Dianne K. Newman

## Abstract

Extracellular electron transfer (EET), the process whereby cells access electron acceptors or donors that reside many cell lengths away, enables metabolic activity by microorganisms, particularly under oxidant-limited conditions that occur in multicellular bacterial biofilms. Although different mechanisms underpin this process in select organisms, a widespread strategy involves extracellular electron shuttles, redox-active metabolites that are secreted and recycled by diverse bacteria. How these shuttles catalyze electron transfer within biofilms without being lost to the environment has been a long-standing question. Here, we show that phenazine electron shuttles mediate efficient EET through interactions with extracellular DNA (eDNA) in *Pseudomonas aeruginosa* biofilms, which are important in nature and disease. Retention of pyocyanin (PYO) and phenazine carboxamide in the biofilm matrix is facilitated by binding to eDNA. In vitro, different phenazines can exchange electrons in the presence or absence of DNA and phenazines can participate directly in redox reactions through DNA; the biofilm eDNA can also support rapid electron transfer between redox active intercalators. Electrochemical measurements of biofilms indicate that retained PYO supports an efficient redox cycle with rapid EET and slow loss from the biofilm. Together, these results establish that eDNA facilitates phenazine metabolic processes in *P. aeruginosa* biofilms, suggesting a model for how extracellular electron shuttles achieve retention and efficient EET in biofilms.

## INTRODUCTION

Microbial biofilms are ubiquitous in natural and engineered contexts, spanning plant roots to chronic human infections to anaerobic digestors (Watnick and Kolter, 2000). As biofilms develop, metabolic stratification occurs, driven by steep concentration gradients of substrates, such as oxygen, that are consumed by cells at the biofilm periphery faster than the substrates can diffuse into the biofilm interior (Stewart, 2003; Stewart and Franklin, 2008; Xu et al., 1998). Indeed, oxidant limitation is a generic challenge for cells that inhabit biofilm microenvironments where electron donors are abundant, yet electron acceptors are not. One widespread strategy microbes employ to overcome this challenge is to channel electrons derived from intracellular metabolism to extracellular oxidants at a distance (Shi et al., 2016). Known as “extracellular electron transfer” (EET), this process requires electron carriers to bridge the gap, be they outer membrane-associated or extracellular cytochromes (Jiménez Otero et al., 2018; Richter et al., 2009; Xu et al., 2018), cytochrome-replete “nanowires” (Subramanian et al., 2018; Wang et al., 2019), cable bacteria conductive filaments (Cornelissen et al., 2018), or redox-active small molecules (Glasser et al., 2017a; Hernandez and Newman, 2001). While the putative molecular components underpinning different EET processes have been described in a variety of organisms, a detailed understanding of how these components achieve EET remains an important research goal across diverse systems.

In contrast to the intense study of microbial nanowires (Malvankar et al., 2011; Reguera et al., 2005; Wang et al., 2019), less attention has been paid to how small soluble (physically diffusive) extracellular electron shuttles facilitate EET beyond interactions at the cell surface (Light et al., 2018; Marsili et al., 2008; Xu et al., 2016). In part, this neglect is due to the challenges involved in identifying and studying small molecule metabolites, compared to the multiheme cytochromes observed in many genomes of organisms known to engage in EET. Accordingly, to study extracellular electron shuttling, we have chosen to work with a model system that employs a relatively well studied and tractable set of shuttles called phenazines. Phenazines are colorful redox-active molecules that are produced by numerous microbial species including the bacterium, *Pseudomonas aeruginosa* (Turner and Messenger, 1986). *P. aeruginosa* strains are ubiquitous yet perhaps most well-known for their roles in chronic infections where their growth as biofilms renders them antibiotic tolerant and contributes to patient morbidity and mortality (Costerton et al., 1999); importantly, phenazines support the development of anoxic, antibiotic tolerant biofilm regions (Dietrich et al., 2013a; Jo et al., 2017; Schiessl et al., 2019). While significant progress has been made in defining the composition of the *P. aeruginosa* biofilm matrix (Colvin et al., 2012) and mapping the zones of phenazine production within it (Bellin et al., 2014, 2016), we still have much to learn about how phenazines facilitate EET within the matrix.

Intriguingly, while the *P. aeruginosa* biofilm matrix comprises a heterogeneous group of polymers, extracellular DNA (eDNA) from dead cells is a significant contributor (Allesen-Holm et al., 2006; Whitchurch et al., 2002), accounting for the majority of the matrix polymers in some cases (Matsukawa and Greenberg, 2004; Steinberger and Holden, 2005). Phenazines have long been known to intercalate into double stranded DNA *in vitro* (Hollstein and Van Gemert, 1971), and more recently, it was suggested that the phenazine pyocyanin (PYO) can participate in DNA-mediated charge transfer (DNA CT) chemistry *in vitro* (Das et al., 2015). Together with the observation that PYO promotes eDNA release by stimulating cell lysis (Das and Manefield, 2012), these facts led to speculation that phenazine-eDNA interactions might facilitate biofilm EET (Das et al., 2015). Notably, the ability of PYO to stimulate cell lysis changes according to the environment: when cells are oxidatively stressed (*i.e.* oxidant replete but reductant limited) and ATP limited, PYO is toxic; whereas when they are reductively stressed (*i.e.* reductant replete but oxidant limited), PYO promotes viability and biofilm aggregate expansion (Costa et al., 2017; Meirelles and Newman, 2018). This observation suggests the intriguing possibility that cell lysis by a small percentage of the population early on might later promote EET once biofilms have developed anoxic zones where extracellular electron shuttles support metabolism. Though a variety of roles for eDNA in biofilms have been proposed, including serving as a structural support, nutrient and/or genetic reservoir (Flemming and Wingender, 2010), to our knowledge, that eDNA may stimulate biofilm metabolism by facilitating EET has not been tested.

The current model of the phenazine redox cycle in biofilms can be broadly defined (Fig. 1A). In anoxic regions, oxidized phenazines are reduced intracellularly by metabolic reactions that support these cells (Glasser et al., 2014, 2017b; Jo et al., 2017; Wang et al., 2010). These reduced phenazines physically diffuse through the extracellular matrix toward the oxic region where they react abiotically with molecular oxygen. Upon re-oxidation, phenazines return to the anoxic region of the biofilm to complete the redox cycle. Although studies have begun to characterize the reactions on either side of the redox cycle, very little is known about how phenazines operate in the intervening extracellular matrix.

**Figure 1.**
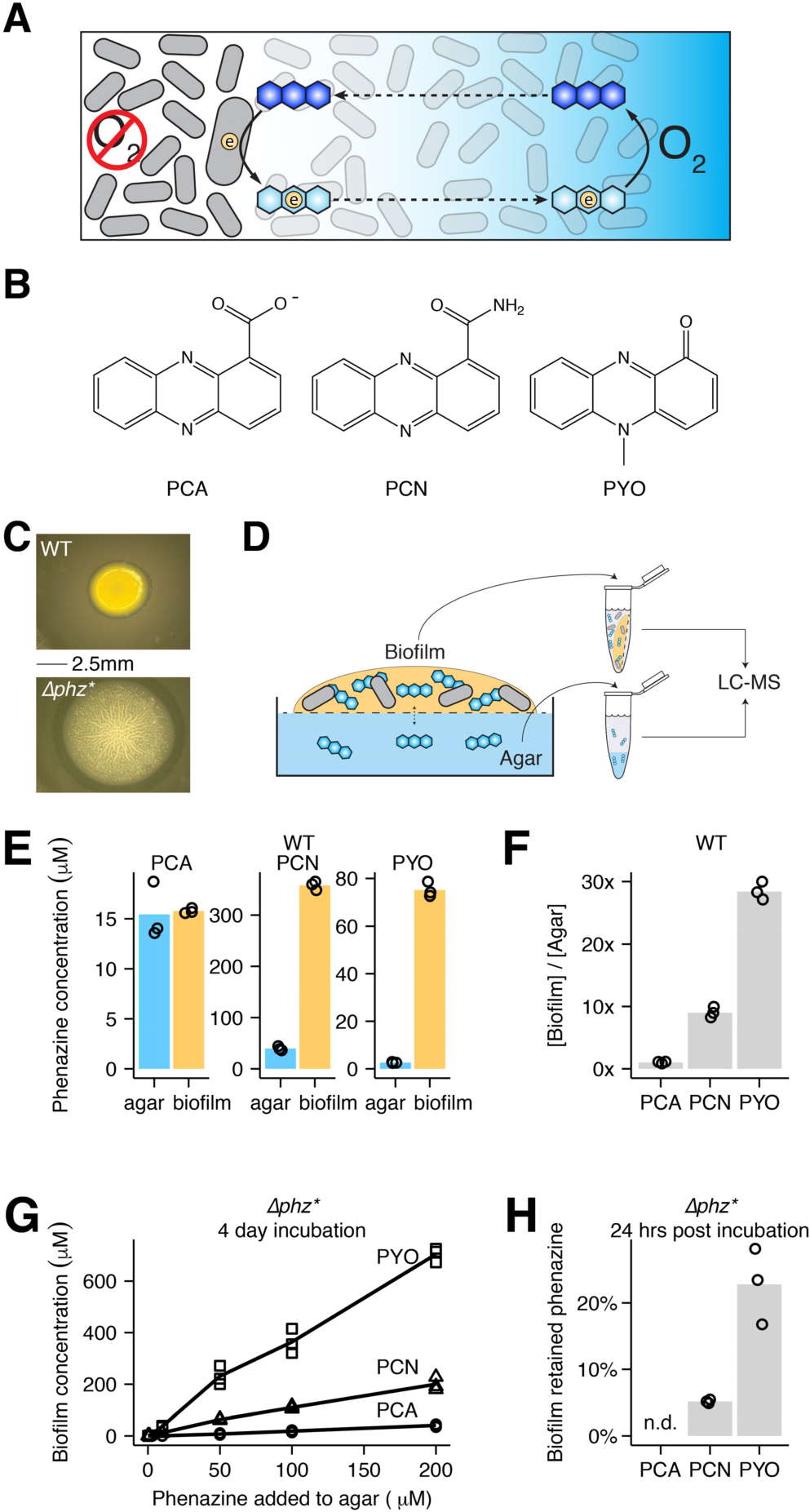
Colony biofilms retain PYO and PCN. (A) Diagram of the phenazine redox cycle in a biofilm. Cells are shown as gray rods, phenazines are shown as blue hexagons, electrons are shown as circles, the oxygen gradient is shown as the blue background. (B) Structures of the three studied phenazines in their oxidized states produced by *P. aeruginosa* – phenazine carboxylate (PCA), phenazine carboxamide (PCN), and pyocyanin (PYO). (C) Images of WT (top) and *Δphz\** (bottom) colony biofilms. (D) Schematic of phenazine extractions from colony biofilms and agar. The 0.2μm membrane is shown as the dashed line. (E) Biofilm and agar concentrations for PCA, PCN and PYO from three WT biofilms. (F) The same data as C, represented as retention ratios ([Biofilm] / [Agar]). (G) Recovered phenazine concentrations from *Δphz*\* colony biofilms grown with different levels of synthetic phenazine in the underlying agar for 4 days. (H) Accumulated phenazine from three *Δphz*\* colony biofilms following three days of growth with synthetic phenazine (Day 3), and one day later after transfer to fresh agar (Day 4). Data are represented as the percentage Day 4 / Day 3. PCA was not detected on Day 4. In panels E-H, values for individual biofilms are shown by open symbols, and lines or bars represent the mean.

Theoretical studies suggest that physical diffusion of oxidized phenazine towards the biofilm interior and reduced phenazine toward the biofilm periphery may be fast enough to support the metabolism of the oxidant limited cells (Glasser et al., 2017a; Kempes et al., 2014). However, these studies assume a closed system, and an unresolved paradox has been how diffusible extracellular molecules could function in a redox cycle without being lost from the biofilm to the environment. Here we explore how phenazine electron transfer may be reconciled with phenazine retention. Specifically, we ask: Are phenazines retained? What mechanisms of electron transfer are compatible with phenazine retention? Is phenazine electron transfer *in vivo* fast compared to phenazine loss? Our motivation to answer these questions arises not only from a desire to constrain the model of phenazine redox cycling within *P. aeruginosa* biofilms, but more broadly, to identify a potentially generalizable strategy for how diverse electron shuttles enable EET.

## RESULTS

We studied three major phenazine derivatives made by *P. aeruginosa* strain UCBPP PA14 (Schroth et al., 2018): phenazine carboxylate (PCA), phenazine carboxamide (PCN), and pyocyanin (PYO) (Fig. 1B). Beyond studying wildtype (WT) produced phenazines, we also explore the effects of individual synthetic phenazines on a mutant that does not produce phenazines, *Δphz* (*ΔphzA1-G1*, *ΔphzA2-G2*), or on a mutant that is also incapable of modifying PCA, *Δphz* (Δphz, ΔphzMS, ΔphzH*). Experiments were performed with two different types of biofilms: macroscopic colony biofilms grown on nutrient agar surfaces (Fig. 1C, Fig. S1A) and microscopic biofilms attached to the surface of an interdigitated microelectrode array suspended in liquid medium. Phenazine-dependent biofilm phenotypes operate similarly at both scales (Ramos et al., 2010), so we selected the biofilm cultivation method for any given experiment based on which was best suited to answering our specific research question.

### Colony biofilms retain PCN and PYO, but not PCA

First, we sought to quantify phenazine retention by colony biofilms (Fig 1C-D). In contrast to previous work that used an electrode array to electrochemically measure the spatial distribution of phenazines that physically diffuse into agar underneath colony biofilms (Bellin et al., 2014, 2016), we used liquid chromatography-mass spectrometry (LC-MS) to quantify extracted endogenous phenazines from the biofilms and compare their concentrations to that in the underlying agar (Fig 1D-F). Colony biofilms could be cleanly separated from the agar, because they were separated by a 0.2µm membrane filter, which did not affect the results (Fig. S1B-C). Overall, PCA, PCN, and PYO concentrations varied by more than 10-fold in the biofilms reaching concentrations of ∼15µM PCA, ∼ 400µM PCN, and ∼80µM PYO. Comparing biofilm to agar concentrations showed that PCN and PYO were enriched in the biofilm 10-fold and 30-fold, respectively, while PCA reached similar concentrations in the biofilm and the agar (Fig. 1E-F). This suggested that PCN and PYO were strongly retained by the biofilm and PCA was not. Importantly, lysing resuspended biofilm cells by sonication prior to phenazine quantification did not strongly affect the results (Fig. S1D), indicating that the measured pools of phenazines were predominantly retained in the extracellular matrix rather than intracellularly.

To test if differential phenazine retention was caused by a spatial or temporal difference in biosynthesis, we grew *Δphz*\* colony biofilms with synthetic phenazines in the agar and quantified phenazines taken up by the biofilm. Incubation with ≥ 10μM PYO resulted in >200µM PYO accumulation in the biofilm (Fig. 1G). PCN accumulated to a lesser extent, and PCA biofilm uptake was minimal (<50µM) even with 200µM added to the agar (Fig. 1G). *Δphz*\* colonies transferred from phenazine agar to fresh agar after 3 days of growth retained phenazines in the same pattern as the wild type (WT) over 24 h (Fig. 1H), demonstrating that the observed phenazine retention does not depend on endogenous phenazine production. Wild-type colony biofilms exhibit relatively thick and smooth morphologies that contain deep anoxic regions that are thought to be supported by phenazines. *Δphz\** colony biofilms exhibit different colony morphologies that are thin and highly wrinkled, which is thought to be a physiological adaptation to maximize surface area and oxygen penetration in the absence of phenazines as shown for *Δphz* (Dietrich et al., 2013b). Notably, only incubation of *Δphz*\* colonies with exogenous PYO appreciably complemented the colony wrinkling phenotype (Fig. S1A). *P. aeruginosa* colony biofilms thus appear able to take up and use significant amounts of exogenous PYO, and PCN to a lesser extent. These results predict that colony biofilms contain an extracellular matrix component that binds and effectively retains PYO and PCN, but not PCA.

### Phenazines differentially bind extracellular DNA

The extracellular matrix in *P. aeruginosa* PA14 biofilms is known to be primarily composed of two polymers: DNA from dead cells (eDNA) and the polysaccharide, Pel (Colvin et al., 2011; Das and Manefield, 2012). To test the hypothesis that eDNA in the biofilm matrix was responsible for binding phenazines, we quantified the binding affinity of oxidized PCA, PCN, and PYO for double-stranded (ds) DNA *in vitro* using isothermal titration calorimetry (Fig. 2A). As expected, oxidized PCA showed no detectable binding because it is negatively charged, as is the phosphate backbone of DNA at pH 7. In contrast, oxidized PCN (K_D_ = 194µM; 95% C.I. 148 – 305µM) and PYO (K_D_ = 13µM; 95% C.I. 6.5 – 49µM) both bind ds DNA, and these results were consistent with ethidium bromide displacement and microscale thermophoresis binding assays (Fig. S2A-B) (Das et al., 2015). Notably, these *in vitro* phenazine-DNA binding affinities correlate with their *in vivo* retention ratio ( [biofilm] / [agar]), where PYO is retained in the biofilm significantly more than PCN, and PCA is not retained. Reduced PYO showed no change in endogenous fluorescence upon addition of calf thymus DNA (Fig. S2C), which is unexpected for strong intercalative binding. Therefore, the DNA binding affinity of PYO is likely redox dependent.

**Figure 2.**
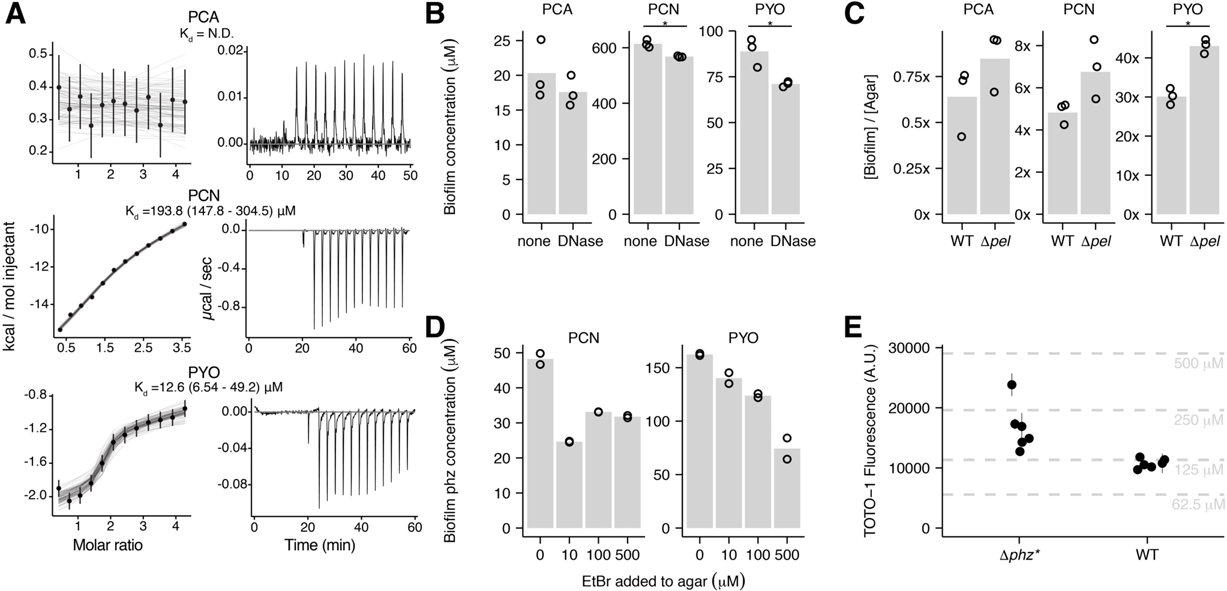
Phenazines interact with DNA *in vitro* and *in vivo*. (A) Representative isothermal titration calorimetry (ITC) data for each phenazine injected into a solution of ds DNA (29bp). Exothermic reactions are depicted as negative values. Integrated peak data was fit with a Bayesian model to calculate the K_d_ (in bp DNA) with 95% confidence intervals (values shown in parentheses) (Duvvuri et al., 2018). (B) Biofilm phenazine concentrations for WT biofilms treated with or without DNase I in the underlying agar for 24hrs (n = 3 per condition). Bars with asterisks denote measurements that differ significantly (p < 0.05) by a Welch’s single tailed t-test. (C) Phenazine retention ratios for WT and *Δpel* colony biofilms (n = 3 per condition). Statistical test same as in B. (D) Accumulated phenazine concentrations for *Δphz\** biofilms incubated with 50uM PCN or PYO and increasing concentrations of the competitive intercalator, ethidium bromide, in the underlying agar. (E) eDNA quantified in six WT and *Δphz\** colony biofilms with the dye TOTO-1. Error bars show standard deviation from two technical replicates. Dashed lines show calf thymus DNA standards, with concentrations back calculated for the biofilm volume. In panels B-D, values for individual biofilms are shown by point symbols, and bars represent the mean.

To determine whether phenazine-eDNA binding occurs *in vivo*, we treated 3-day old WT biofilms with DNase I for 24 hrs. These experiments were performed with DNase I spotted on tryptone agar medium rather than its optimal buffer, as controls showed that buffer alone significantly disturbed the biofilm (Fig. S3A-C). Despite a low activity for DNase under these conditions, DNase-treated biofilms showed significantly lower biofilm PCN and PYO concentrations than their untreated counterparts; moreover, PCA concentration was unchanged (Fig. 2B). *P. aeruginosa* eDNA originates from the genomic DNA of dead cells and is therefore high molecular weight and may be bound by other biomolecules (e.g. proteins) (Kavanaugh et al., 2019). Therefore, DNase treatment was likely only partially effective because it could not cleave ds DNA in the presence of other bound matrix components, and/or because it did not have enough activity to degrade the eDNA completely to eliminate phenazine binding sites. We also compared the phenazine retention in the WT to a Pel mutant *(Δpel*) and found that biofilms without Pel retained significantly more PYO (Fig. 2C). Because Pel is known to bind eDNA (Jennings et al., 2015), these results suggest that Pel may partially block access to eDNA by PYO, although this remains to be tested *in vitro*.

To probe the eDNA binding sites within the biofilm using a different approach, we competed phenazines against ethidium bromide, a classical DNA intercalator. Since PCN and PYO compete for DNA binding sites with ethidium *in vitro*, and ethidium is largely excluded from cells (Jernaes and Steen, 1994), we reasoned that these intercalators could compete for binding sites in the biofilm eDNA. We grew *Δphz\** biofilms with 50µM PCN and PYO and increasing amounts of ethidium in the underlying agar. Figure 2D shows that increasing concentrations of ethidium resulted in successively less PYO accumulating in the biofilms, while PCN accumulated to a similar lower level in the presence of any amount of added ethidium.

Confocal microscopy of WT and *Δphz\** colony biofilms with a cell-impermeable ds DNA dye, TOTO-1 (Okshevsky and Meyer, 2014), showed abundant eDNA localized in dead cells and in between cells (Fig. S3D). We quantified the bulk concentration of eDNA in colony biofilms by incubating biofilm suspensions with TOTO-1 and measuring dye fluorescence. Both WT and *Δphz\** biofilm suspensions yielded large fluorescence values when incubated with TOTO-1. These values fall within the range of 60 – 500µM bp ds DNA in the colony biofilms, when calibrated against standards of calf thymus DNA (Fig. 2E). However, adding calf thymus DNA to the biofilm suspensions did not yield the expected increase in dye fluorescence (Fig. S3E), which suggests that the dye may be partially inhibited by biofilm components. Therefore, this order of magnitude estimate of biofilm eDNA should be interpreted as a lower bound on the true value. Given this estimate, the biofilm eDNA (>100µM bp) is in excess of PYO (∼80µM), but it may not be in excess of PCN (>300µM). Due to its poor aqueous solubility, it is probable that PCN crystallizes extracellularly at the observed biofilm concentrations, which could lead to its measured retention (Hernandez et al., 2004). Together, our *in vivo* and *in vitro* results are consistent with eDNA providing binding sites for oxidized PCN and oxidized PYO in the biofilm extracellular matrix.

### Constraints on phenazine electron transfer mechanisms *in vitro* and *in vivo*

Given that phenazines are differentially bound and retained in biofilm eDNA, we next sought to constrain how electron transfer might be achieved in this context. Previous research has shown distinct localization patterns for different phenazines within biofilms, with the lowest potential phenazines (*e.g.* PCA) in the interior, and the highest potential phenazine (*e.g.* PYO) at the oxic periphery (Bellin et al., 2014, 2016). To test whether electron transfer could occur between these molecules in solution, we mixed different oxidized and reduced phenazines under anoxic conditions and monitored the absorbance spectra before and after mixing (Fig. 3A-B). Because PYO exhibited the largest changes in absorbance upon reduction, we monitored different mixtures of PYO with PCA or PCN at 690nm (unique PYO absorbance maximum) starting one minute after mixing, at which point equilibrium had been achieved.

**Figure 3.**
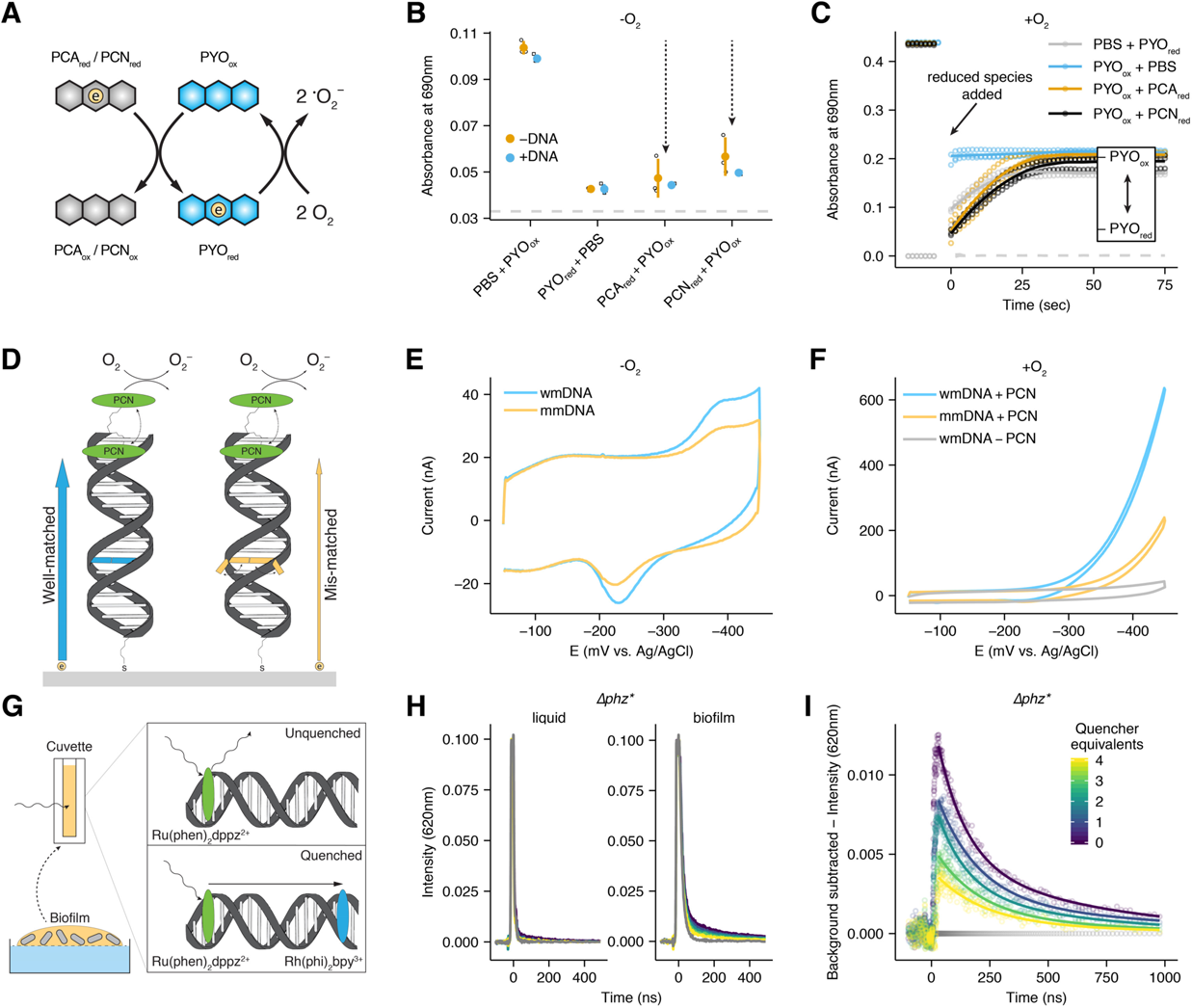
Inter-phenazine electron transfer and DNA CT. (A) Diagram showing an electron transfer reaction in solution between a reduced phenazine and oxidized PYO and between reduced PYO and molecular oxygen. (B) Reaction progress after 1 min measured at 690nm for mixtures of phenazines shown in A, compared to oxidized and reduced PYO alone. Each reaction was performed in the presence and absence of calf thymus DNA. For each condition n=3 and error bars are one standard deviation. Dashed line shows the background signal from PBS alone or with PCN or PCA. (C) PYO oxidation state measured at 690nm over time (diagnostic for oxidized PYO) for different reactions in the presence of oxygen. Points are individual measurements, lines are loess smoothed for each set of triplicate measurements. Dashed line shows the background signal from PBS with PCN_red_ or PCA_red_. (D) Schematic showing a DNA modified electrode with tethered PCN (green oval) and the expected electron transfer for well-matched duplexes (blue arrow and bp) and duplexes containing a mismatch (orange arrow and bp). Mismatched bases are less likely to be in a well stacked position, which is necessary for electron transfer through the DNA p-stack. (E) Representative cyclic voltammetry of the well-matched (wmDNA) and mismatched (mmDNA) constructs shown in D under anoxic conditions. (F) Representative cyclic voltammetry of the well matched, mismatched, or no phenazine constructs under the aerobic conditions described in D. (G) Diagram of time resolved spectroscopy of the photoexcited electron donor Ru(phen)_2_dppz^2+^ quenched by Rh(phi)_2_bpy^3+^ with biofilm eDNA. (H) Comparison of Ru(phen)_2_dppz^2+^ fluorescence in the presence of a concentrated liquid *P. aeruginosa* culture and a resuspended biofilm containing eDNA. Gray lines show background biological fluorescence before Ru(phen)_2_dppz^2+^ was added. The color map is the same as I. (I) The background subtracted data from the biofilm panel of H. The amount of Rh(phi)_2_bpy^3+^ is color coded as quencher equivalents relative to Ru(phen)_2_dppz^2+^. Dots are raw data, lines are fit biexponential decays.

Reactions proceeded as expected from the redox potentials of the phenazines, where PYO was almost completely reduced by the lower potential PCA and PCN, but reduction of PCA and PCN by the higher potential PYO was minimal (Fig. 3B, Fig. S4A). In addition to establishing that electron transfer can occur between different phenazines, given their similar structures, these results suggest that electron transfer between like phenazines (*e.g.,* PYO-PYO electron self-exchange) can occur within a redox gradient. Moreover, reactions between reduced PCA or PCN and oxidized PYO proceeded faster than oxidation of any of these phenazines by molecular oxygen (Fig. 3C). We next wondered whether the presence of eDNA would affect the extent of PYO reduction. PYO reduction by PCA or PCN proceeded to completion in the presence of eDNA (Fig. 3B). Because PCA does not bind eDNA, this result suggests electron transfer is occurring in solution between PCA and unbound PYO. For PCN and PYO that both bind eDNA, it is also possible that electron transfer is achieved by their unbound counterparts in solution. However, it has long been known that DNA can facilitate electron transfer between bound redox molecules (Genereux and Barton, 2010), motivating us to test whether such a process could also occur within our *P. aeruginosa* biofilms.

DNA facilitates charge transfer (DNA CT) through the π-stacked base pairs (Genereux and Barton, 2010), and recent studies have shown that DNA CT can occur over kilobase distances (Tse et al., 2019). A previous study suggested that PYO might be able to transport electrons via DNA (Das et al., 2015), but given the preliminary nature of these experiments, we decided to revisit these experiments more rigorously. To better test the ability of phenazines to carry out DNA CT, we covalently attached a phenazine via a flexible linker to one DNA strand and then made DNA-modified gold electrodes with a thiol linker on the complementary strand according to standard protocols (Kelley et al., 1997; Slinker et al., 2010, 2011). Specifically, short ds DNA molecules (17 bp) were covalently linked to the gold surface to form a packed monolayer, and the distal 5’ end of each duplex contained a covalently linked PCN, the phenazine derivative most amenable to synthesis (see Materials and Methods) (Fig. 3D). Thus, we established a well-defined chemical system to test if a phenazine could participate in electron transfer to the electrode through the ds DNA.

Because the efficiency of DNA CT depends upon the integrity of the π-stacking of base pairs within the DNA duplex (Genereux and Barton, 2010), we compared well-matched duplex DNA monolayers to duplex DNAs containing a single base mismatch that stacks less efficiently (Fig. 3D). We utilized multiplexed DNA chips to facilitate replicate comparisons between well-matched and mismatched DNA monolayers (Fig. S4B-D); measurements with a non-intercalating control probe showed that these different monolayers had very similar surface coverages (Fig. S4E-F) (Slinker et al., 2010). Figure 3E shows that the mismatched construct yielded diminished current in the phenazine redox peak, consistent with the charge transfer being DNA-mediated; the presence of the intervening mismatch inhibits DNA CT. This mismatch effect was consistent across replicate low density (32-54% decrease) and high density (36-69% decrease) DNA monolayers (Fig. S4C and supp. text). These results displayed mismatch attenuation similar to that observed for well characterized small molecules shown to stack with the DNA duplex and carry out DNA-mediated CT (Slinker et al., 2011). Strikingly, in the presence of oxygen, a classic voltammetric signal characteristic of electrocatalysis was obtained (Fig. 3F) centered on the phenazine redox peak. Hence the DNA-tethered phenazine is able to accept electrons from the electrode through the DNA π-stack and then reduce oxygen in a catalytic fashion. Together, these results demonstrate that a *P. aeruginosa* phenazine can participate in DNA CT *in vitro*.

To test if biofilm eDNA could support DNA CT, we incubated *Δphz\** colony biofilms suspended in PBS with well-characterized intercalators that can perform DNA CT reactions, which can be monitored by time-resolved spectroscopy (Arkin et al., 1996a). The quenching of photoexcited Ru(phen)_2_dppz^2+^ by Rh(phi)_2_bpy^3+^ is well-characterized and known to occur by a redox mechanism (Stemp et al., 1995) and not energy transfer (e.g. FRET), with both the forward and back electron transfers occurring predominantly on the picosecond timescale (Fig. 3G, Fig. S4G) (Arkin et al., 1996b). Both of these intercalators bind to ds DNA more than an order of magnitude more tightly than do phenazines.

Moreover, Ru(phen)_2_dppz^2+^ is luminescent in aqueous solution only when intercalated in DNA (or otherwise protected from water), precluding a 2^nd^ order reaction between the complexes in solution. Thus, in time-resolved emission experiments on the nanosecond timescale, static quenching, where quenching is fast and occurs without a change in the Ru(phen)_2_dppz^2+^ excited state lifetime, is consistent with DNA-mediated CT (Arkin et al., 1996a), while slower dynamic quenching, which leads to a change in emission lifetime, is consistent with a slower diffusive process. We first compared the Ru(phen)_2_dppz^2+^ signals of liquid grown and biofilm suspensions (of the same optical density) to determine if the signal was specific to eDNA; the ruthenium complex is not expected to be taken up by the cells and bind to genomic DNA on the time scale of this experiment (Fig. 3H). We only observed ruthenium luminescence in the presence of the biofilm suspension, consistent with ruthenium luminescence being associated with binding to eDNA. We then examined the pattern of quenching of Ru(phen)_2_dppz^2+^ by Rh(phi)_2_bpy^3+^. Figure 3I shows static quenching where the intensity of the Ru(phen)_2_dppz^2+^ signal decreases, while the observed decay kinetics are unchanged (Fig. S4H). Therefore, we conclude that biofilm eDNA can support rapid DNA CT between these two metal complexes, faster than the timescale for diffusion.

### Electrode-grown biofilms retain PYO capable of extracellular electron transfer

Having explored phenazine retention and electron transfer separately, we next wanted to establish a system in which we could monitor both of these processes simultaneously *in vivo*. We took an electrochemical approach to achieve this, growing *P. aeruginosa* biofilms on interdigitated microelectrode electrode arrays (IDA) (Fig. 4A). Biofilms were grown by incubating IDAs in bioelectrochemical reactors with planktonic cultures under oxic conditions and stirring (Fig. 4B). After 4 days, with medium replaced daily, mature biofilms were transferred to anoxic reactors with fresh medium for electrochemical measurements. Confocal and SEM imaging revealed that the IDA biofilms were heterogeneous in composition, but consistently contained multicellular structures of live cells and abundant eDNA (Fig. 4C-E, Fig. S5A-B). Under these conditions, the cells predominately produced PYO, as measured by LC-MS (Fig. S5C). Because PYO was the most tightly retained phenazine in colony biofilms and *in vitro*, we focused on this phenazine for the remainder of these experiments.

**Figure 4.**
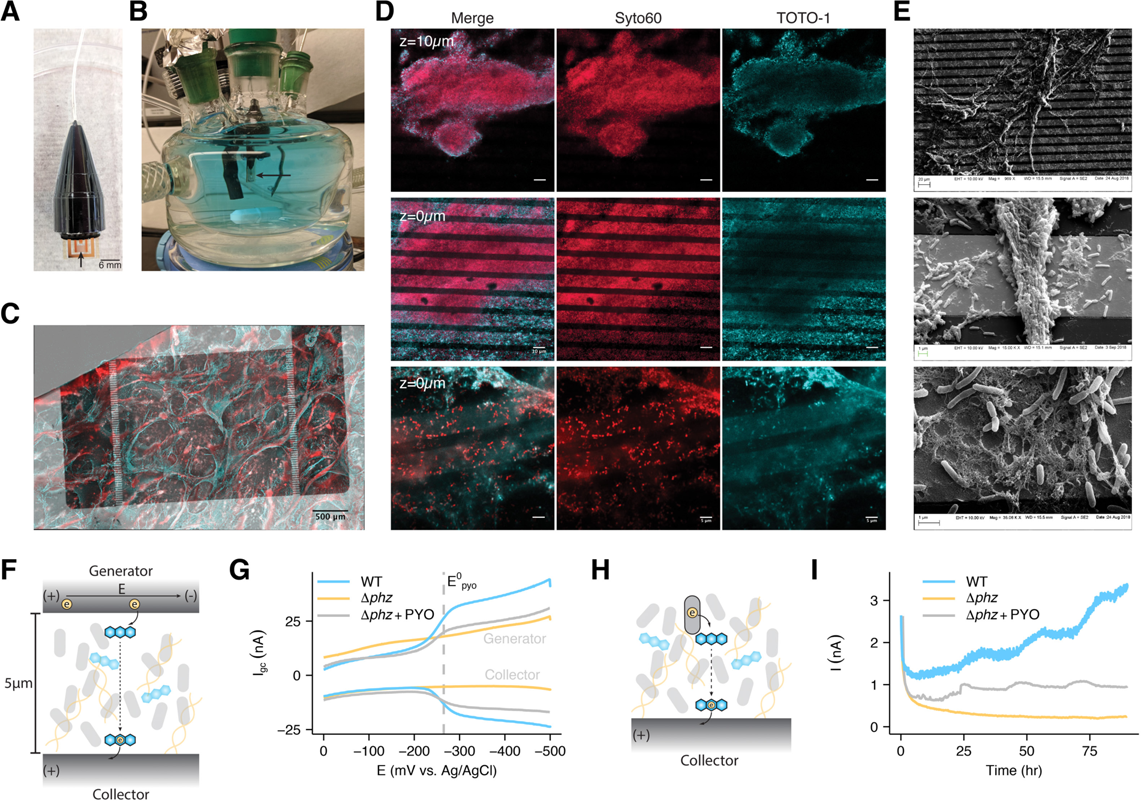
*P. aeruginosa* forms biofilms on electrodes and shows PYO dependent conductivity. (A) Photograph of a sterile IDA. (B) Photograph of the growth/electrochemical reactor with a submerged IDA in a fresh medium + PYO solution. (C) A max intensity projection of a WT IDA biofilm imaged using Syto60 (cell permeable – all DNA), shown as red, and TOTO-1 (cell impermeable – eDNA), shown as cyan. Fluorescence channels are overlaid on a transmitted light channel showing the opaque gold regions of the electrode in gray scale. (D) Fluorescence microscopy of IDA biofilms with the same dyes as in C. Top: a 63x confocal image of a *Δphz*\* IDA biofilm from a zstack 10μm above the electrode surface (scale bar = 10μm). Middle: a confocal image from the same zstack as top, but at the electrode surface (scale bar = 10μm). Bottom: a confocal slice of a different *Δphz*\* biofilm from a 63x zoom with Airyscan, showing single live cells and eDNA on the electrode surface (scale bar = 5μm). (E) SEM images at increasing magnification showing cells and extracellular matrix on the IDA electrode bands. (F) Diagram showing how a generator-collector (GC) two electrode system can generate current through PYO reduction. (G) GC data is displayed as the current at each electrode vs. the generator potential, E. Representative measurements are shown for WT, *Δphz* and *Δphz* + PYO biofilms. (H) Diagram showing how cells generate metabolism dependent current through phenazine reduction. (I) Metabolic current described in H is measured by chronoamperometry for *Δphz* and WT biofilms over several days.

Originally used to measure conductivity of abiotic materials, the IDA has a 2-working electrode geometry and recently was adapted to study EET through microbial biofilms (Boyd et al., 2015; Snider et al., 2012; Xu et al., 2018; Yates et al., 2015). Measurements are made by driving electron transfer between the two electrode bands across a 5µm gap (Fig. 4F), which treats the biofilm like an abiotic material, resulting in EET that is decoupled from the cells’ metabolic activity. Specifically, we used a generator-collector (GC) strategy to measure EET through the biofilm, where the “generator” electrode is swept from an oxidizing potential (E = 0mV vs. Ag/AgCl) to a reducing potential (E = −500mV), while the “collector” electrode maintains a fixed oxidizing potential (E = 0mV). In this GC arrangement, electron transfer into the biofilm from the generator occurs as the potential of the generator is swept negatively, reducing PYO (E^0^ = −250mV) at the biofilm/generator interface. Electrons are conducted across the gap through the biofilm due to EET, either by physical diffusion of PYO or other mechanisms. The electron transfer at both the generator and collector is measured as current (I_gc_) and plotted against the potential of the generator electrode. Implicit in generating a sustained I_gc_ is recycling of the redox molecules that support the observed current. Figure 4G shows that WT and *Δphz*\* + PYO biofilms supported current (above background) across the 5µm gap over a ∼3 min scan as the generator potential approached PYO’s redox potential (−250mV vs. Ag/AgCl), consistent with PYO-mediated EET; significantly, *Δphz*\* alone did not. Dependency of current on the generator potential observed here is consistent with PYO-mediated EET being a diffusive process. Reduction of PYO at the generator and oxidation of PYO at the collector generates a redox gradient that drives EET through the intervening biofilm, resulting in current centered on the PYO midpoint potential that saturates at strongly reducing generator potentials (Snider et al., 2012). Here the diffusive nature of EET would reflect either physical diffusion of PYO through the biofilm (reduced PYO from the generator to collector, oxidized PYO from the collector to the generator) or the effective diffusion of electrons through bound PYO (Strycharz-Glaven et al., 2011a).

To test whether these short-term GC measurements of biofilm EET indicate long-term ability to support metabolic activity, we poised the IDA electrodes at an oxidizing potential (+100mV vs. Ag/AgCl) and monitored the current produced by the biofilm over 4 days in the presence of 40 mM succinate as the electron donor for cellular metabolism under anoxic conditions. In this configuration the observed current would be due to cellular oxidation of succinate coupled with PYO-catalyzed EET to the poised electrode, where the electrode acts as the terminal electron acceptor instead of oxygen (Fig. 4H). Figure 4I shows that both WT biofilms relying solely on endogenous PYO and *Δphz*\* biofilms + exogenous PYO generate robust current over many days, while *Δphz*\* alone does not. The daily periodic rise in current likely reflects the impact of slight temperature or light fluctuations in the room on the cells’ metabolic activity over the course of the experiment (Kahl et al., 2016, 2019). Differences in current magnitude between the WT and *Δphz*\* + PYO runs are likely due to differences in PYO abundance in the biofilms and/or efficiency of cellular PYO reduction. Together, these results demonstrate that the IDA biofilms can use retained PYO for EET to support metabolic activity. To determine whether our IDA biofilms retained phenazines in the same manner as colony biofilms, *Δphz*\* IDA biofilms were soaked in PYO in one reactor and then transferred to another reactor with fresh medium lacking PYO (Fig. 5A). The equilibration of PYO (from the IDA biofilm) with the fresh medium was monitored by square wave voltammetry (SWV) (Fig. 5B), for which peak current (I_swv_) is proportional to the concentration of the PYO remaining in the biofilm at each time interval (Bard et al., 1980). Thus, the rate of decay of I_swv_ provides a means to assess the loss rate of PYO from the biofilm, a measure of how tightly PYO is retained. Upon transfer, the biofilm PYO peak current (I_swv_) decays in 30-45 min, while for a blank IDA (no biofilm) dipped in PYO, I_swv_ decays in 2-3 min (Fig. 5E). We compared PYO and PCA soaks and found that PCA immediately became undetectable by SWV or GC in the transfer reactor (Fig. S5D-E) because it quickly diffuses out of the biofilm, as expected because it does not bind eDNA. Thus, like colony biofilms, IDA biofilms appear to tightly retain PYO but not PCA.

**Figure 5.**
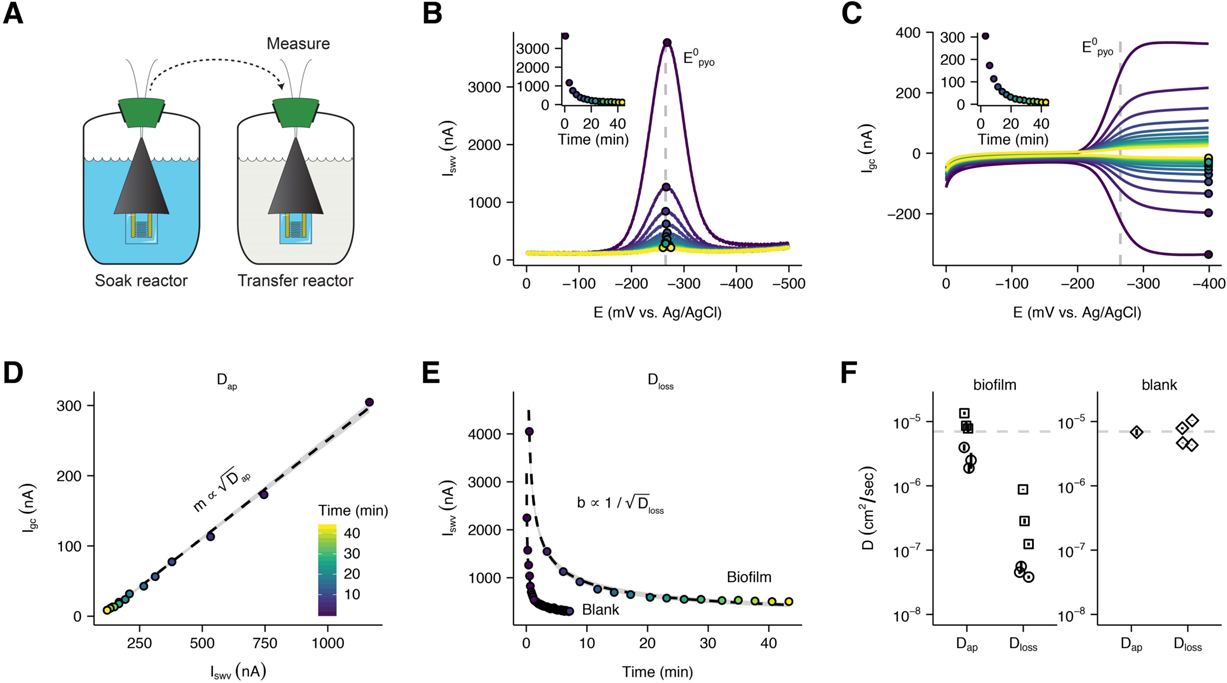
PYO mediated electron transfer is faster than PYO loss. (A) Schematic showing how measurements are made to maximize biofilm specific PYO signals, by transferring IDAs to fresh medium. PYO is shown in blue. (B) Repeated SWV measurements were taken over time as the *Δphz*\* + PYO biofilm equilibrates with the solution in the transfer reactor. Points show peak current within each scan and inset shows the quantified peak currents vs. time. The color map is the same for panels B-E. (C) Repeated GC measurements were taken concurrently with SWV measurements over time as the same biofilm equilibrates. Points and inset are same as in B. (D) Plot of the peak GC vs. peak SWV currents fit with a line (gray area is 95% confidence interval). (E) Comparison of SWV signal over time in the transfer reactor from IDAs with or without a biofilm. Data are fit with a 1D diffusion model, as discussed in the supplementary text. Dashed lines are best fit models and gray regions show 95% confidence intervals. (F) Measurements of D_ap_ and D_loss_ for two *Δphz*\* biofilm IDAs soaked in 75μM PYO (shown as sets of open circles and squares) and a blank IDA soaked in different concentrations of PYO (open diamonds). A parameter for calculating D_loss_ (scan time - t_s_) was solved by assuming that D_ap_ = D_loss_ for the blank IDA (see supplementary text). Error bars for D_ap_ are 95% confidence intervals from the linear fit. Error bars for D_loss_ assume *I_0_* was the peak current in the soak reactor and high and low estimates were generated with the 95% confidence interval values from the SWV fit as well as the D_ap_ estimate. The dotted line is at 7 x 10^-6^ cm^2^ / sec, the measured D for the similarly sized small molecule, caffeine (Niesner and Heintz, 2000).

### Electron transfer through biofilms is faster than PYO loss

Next, we sought to understand the efficiency of PYO-mediated EET in the biofilm. We reasoned that a determination of “efficiency” would compare the rate of electron transfer (which supports the metabolism of the O_2_ limited cells) to the loss rate of PYO from the biofilm (which limits the utility of each PYO molecule). These two processes can both be described by diffusion coefficients, so in a single electrochemical experiment we measured the apparent diffusion coefficient for the PYO-mediated EET, D_ap_, which characterizes all of the redox processes with the electrode, and the diffusion coefficient for PYO as it is lost from the biofilm, D_loss_.

We determined D_ap_ for EET by PYO in an IDA *Δphz*\* biofilm to avoid confounding PYO retention with production, although WT biofilms with endogenous PYO yielded similar results (Fig. S6A-B). Each *Δphz*\* biofilm was soaked with PYO in one reactor then transferred to a second reactor lacking PYO (Fig. 5A), and the equilibration of the biofilm PYO into the fresh medium was monitored by paired SWV and GC measurements over time (Fig. 5B and Fig. 5C). Noting that I_swv_ is proportional to 𝐶 ∗ 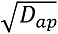, where C is the effective PYO concentration (Bard et al., 1980), whereas I_gc_ is proportional to 𝐶 ∗ 𝐷_ap_ (Strycharz-Glaven et al., 2011b), plotting I_gc_ vs. I_swv_ for each time point is expected to yield a linear dependency with a slope (*m*) proportional to 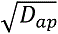 (Fig. 5D) when D_ap_ is independent of the concentration of PYO in the biofilm (Akhoury et al., 2013; White et al., 1982a). In this manner, it is possible to measure D_ap_ for PYO remaining in the biofilm at any given instance when its concentration is unknown.

Applying this approach, for two biological replicates of *Δphz*\* biofilms (three technical replicates each), we found that the mean D_ap_ for PYO is 6.4 x10^-6^ cm^2^/sec over a wide range of PYO biofilm concentrations (Fig. 5F, Fig. S6C). To validate our approach, we measured D_ap_ using a blank IDA with no biofilm, only solution PYO for which we expect D_ap_ ≍ D_loss_. Using known PYO concentrations with a blank IDA (no biofilm) we obtained nearly identical estimates (D_ap_ ≍ 6.8×10^-6^ cm^2^/sec) using our I_gc_ vs. I_swv_ method (concentration unknown) or established methods (I_gc_ vs. [PYO] and I_swv_ vs. [PYO]) that depend on known concentrations (Fig. 5F, Supplementary text, Fig. S6D-G) (Bard et al., 1980). We further validated our measurement scheme by comparing it to an established chronocoulometry technique with two other redox molecules (Hexaammineruthenium(III) and ferrocenemethanol) in the presence and absence of the polymer Nafion. In all cases estimates of D_ap_ by the two methods were within ∼2x (Fig. S6H).

To estimate the upper limit for D_loss_ of PYO lost from *P. aeruginosa* biofilms, we applied a semi-infinite one-dimensional diffusion model (Supplemental Text) to fit the decay of the same I_swv_ measurements used above (Fig. 5B, 5E). Although each SWV scan depends on D_ap_, the decay process of I_swv_ results from loss of PYO out of the biofilm (D_loss_). This calculation requires a scan time constant, whose value we constrain by using the blank control where we assume D_ap_ = D_loss_, allowing us to solve for this constant (see Supplemental Text and Fig. S7A). For *Δphz*\* biofilms, this diffusion model yields a mean D_loss_ of 2.0×10^-7^ cm^2^/sec (Fig. S7B). Hence, D_ap_ for PYO-mediated biofilm EET is more than 25-fold higher than D_loss_ (Fig. 5F). While this model simplifies the actual physical system, it provides a means to estimate an upper limit for the rate of PYO loss from the biofilm. Such a low D_loss_ is consistent with the relatively long time it takes for I_swv_ to decay when transferred to fresh medium for PYO in a biofilm (∼45min) compared to PYO for a blank IDA (<2min) (Fig. 5E) or to PCA in a biofilm (< 1 min) (Fig. S5E). Moreover, the conservative assumptions of our model make it likely that we have underestimated the true difference between D_loss_ and D_ap_ in the biofilm. Collectively, these observations support the idea that PYO EET occurs rapidly compared to the loss of PYO diffusing out of the IDA biofilms.

## DISCUSSION

The redox activity of phenazine metabolites produced by *P. aeruginosa* has intrigued researchers since the 1930’s (Friedheim, 1934), and over the last twenty years a model has emerged that links phenazine electron transfer to biofilm metabolism (Dietrich et al., 2013b; Hernandez and Newman, 2001; Jo et al., 2017). Using the well-characterized *P. aeruginosa*-phenazine system as a model for studying EET mediated by extracellular electron shuttles, here we addressed the conundrum of how phenazines complete their redox cycle within the biofilm matrix without being lost to the environment. Our results point to eDNA as being a critical component of the matrix that facilitates phenazine cycling for EET.

Quantifying phenazine retention in a simple biofilm system was our first goal. While past work has measured phenazines in culture supernatants and in agar underlying colony biofilms (Bellin et al., 2014, 2016; Dietrich et al., 2006), phenazine concentrations within any type of biofilm system were unknown. We found that the degree of retention varied dramatically between the three studied phenazines in colony biofilms, with PCN and PYO being strongly enriched in the biofilm, in contrast to PCA, which readily diffuses away. We observed a similar trend for biofilms grown in liquid medium attaching to IDAs. Notably, oxidized PCN and PYO bind ds DNA *in vitro*, and perturbing extracellular DNA binding sites disrupted PYO retention (and PCN to a lesser extent) *in vivo*. Previous studies have shown PYO actually promotes eDNA release via cell lysis, so PYO retention by eDNA suggests a connection between eDNA production and utilization. To our knowledge, PYO retention in eDNA is the first example of a metabolically helpful (as opposed to biofilm structural) molecule being bound by the extracellular matrix of a biofilm. Because eDNA is found in biofilms made by diverse species and many small molecules exhibit DNA binding capacity, our results may be broadly relevant to diverse biofilm functions involving extracellular metabolites.

Recognizing that a primary function for phenazines is extracellular electron transfer, we characterized this process *in vivo*. Our IDA experiments suggest that PYO simultaneously can be retained (low D_loss_) and facilitate fast EET (high D_ap_), establishing the efficiency of this process. To understand the mechanism that underpins this efficient EET, we can interpret our results in the context of electron transfer theory from the study of redox polymers (Dalton and Murray, 1990). This theory holds that the measured D_ap_ is the sum of the effective diffusion coefficient of electrons due to self-exchange reactions among bound shuttles (D_e_) and the physical diffusion coefficient (D_phys_) of any unbound shuttles (D_ap_ = D_e_ + D_phys_). In this context, we think there are two ways to explain our electrochemical results that biofilm PYO D_ap_ (∼6.4 x10^-6^ cm^2^/sec) is higher than biofilm PYO D_loss_ (∼2.0 x10^-7^ cm^2^/sec), and similar to solution PYO D_phys_ (6.8 x10^-6^ cm^2^/sec) (Fig. 5F). First, if we assume our measured D_loss_ is the same as PYO D_phys_ within the biofilm, the difference between D_ap_ and D_phys_ can be explained by D_e_. Our *in vitro* data suggest that such self-exchange reactions (D_e_) could be DNA-mediated. In the case of very rapid electron transfer among eDNA-bound PYO, D_ap_ would still be limited by counter ion diffusion, consistent with a measured D_ap_ similar to that of a small molecule in solution (∼7 x 10^-6^ cm^2^/s) (White et al., 1982b). Alternatively, the measured loss of PYO from the biofilm, D_loss_, may not accurately reflect physical diffusion of PYO within the biofilm, D_phys_. Because the IDA biofilms do not completely cover the electrodes, PYO may be able to physically diffuse in solution adjacent to them. In this case, diffusion in solution would be consistent with the measured D_ap_, and the low D_loss_ measurement would reflect the slow release of PYO from the IDA biofilm due to its retention by eDNA. Regardless, the striking measured difference between D_ap_ and D_loss_ indicates that PYO electron transfer promotes efficient biofilm EET metabolism because electron transfer occurs rapidly, while loss of PYO to the environment is slow.

Together, our results allow us to refine our model for how phenazine EET may operate within biofilms (Fig. 6). Intriguingly, PYO biosynthesis requires O_2_, whereas PCN and PCA do not, and electrochemical imaging underneath colony biofilms has shown lower potential phenazines in the anoxic interior (PCA, PCN) and the higher potential phenazine (PYO) near the oxic periphery (Bellin et al., 2014, 2016). Reduced PYO is also known to react with oxygen significantly faster than PCA and PCN (Wang and Newman, 2008). As such, it is tempting to speculate that phenazines are ordered in the biofilm matrix in a sequence akin to a large extracellular electron transport chain—from reduced PCA/PCN in the anoxic interior, to eDNA bound PYO at the oxic periphery, and ultimately to molecular oxygen (Fig. 6A). How then do phenazines exchange electrons within this framework? Noting that the heterogeneity of the biofilm matrix makes it possible that different electron transfer pathways could occur in these complex systems, we favor two mechanisms mediated by eDNA for how phenazine EET may operate in the matrix (Fig. 6B). Both mechanisms assume that reduced PCA and PCN will diffuse from the anoxic zone to the oxic zone (i), where PYO_ox_ is bound to eDNA. In one model (Fig. 6B top), PYO’s binding equilibrium will result in some PYO_ox_ unbinding from the eDNA, allowing it to react with PCA_red_ or PCN_red_ (ii). PYO _red_ then reacts with oxygen generating PYO_ox_ (iii), which rebinds DNA. In the other model (Fig. 6B bottom), reduced phenazines (likely PCN) intercalate into eDNA and reduce PYO_ox_ via DNA CT (ii). PYO _red_ unbinds DNA, reacts with oxygen (iii), and PYO_ox_ rebinds DNA. Given that PCA_red_ and PCN_red_ react more quickly with PYO_ox_ than O_2_, then both models would facilitate the re-oxidation of the interior phenazines and promote diffusion back toward the anoxic interior (iv). These non-mutually exclusive models integrate what is known about phenazine O_2_ reactivities, redox potentials and biosynthesis zones, and help explain how PYO and eDNA interactions enhance EET in *P. aeruginosa* biofilms by facilitating retention and electron transfer. Testing these models in physicochemically well-defined matrixes in addition to complex biofilms represents an exciting challenge for future research.

**Figure 6.**
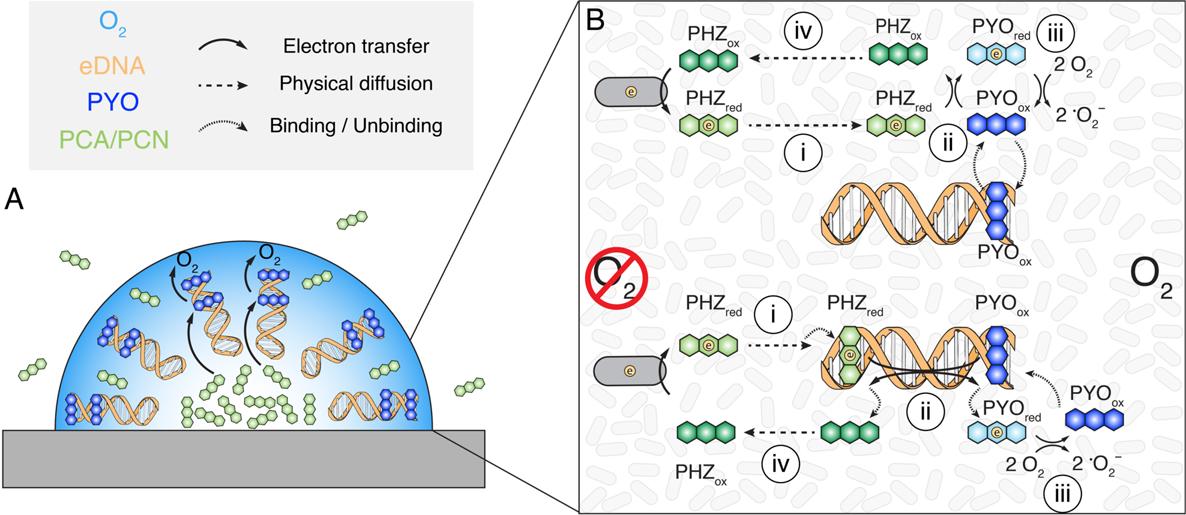
Proposed models of phenazine electron transfer and retention. (A) Overview of phenazine electron transport to oxygen in the biofilm. Note the DNA molecules are drawn radially for explanatory purposes only. (B) Proposed models for how the scheme shown in A occurs. The top model shows electron transfer in solution and the bottom model shows electron transfer via DNA CT. In both models oxidized PYO is mostly bound and retained in the oxic region of the biofilm. PCN and PCA (PHZ) are reduced in the anoxic region of the biofilm and diffuse outward (i). The reduced phenazine reduces oxidized PYO (ii). Reduced PYO then reacts with molecular oxygen (iii), and the oxidized PYO re-equilibrates with the DNA binding sites. This facilitates the re-oxidation of PCA and PCN, allowing some diffusion back toward the interior (iv).

In conclusion, our findings illustrate that eDNA binding provides a mechanism to resolve how otherwise diffusible extracellular electron shuttles can catalyze efficient EET in real world, open systems. Beyond serving as a structural support, carbon source, or genetic reservoir, our studies reveal that interactions between extracellular electron shuttles and eDNA in biofilms underpin their metabolic vitality. It is noteworthy that eDNA is abundant in many biofilms (Flemming and Wingender, 2010) and diverse biofilm-forming bacteria have the potential to produce extracellular electron shuttles (Glasser et al., 2017a). Accordingly, eDNA retention of these electron shuttles—and perhaps of other biologically useful molecules—may represent a widespread strategy whereby a reactive extracellular matrix supports bacterial biofilms in unexpected and physiologically significant ways.

## Acknowledgments

We thank Jeanyoung Jo, Lars Dietrich and Matthew Parsek for providing strains. This work was supported by grants to D.K.N. from NIH (1R01AI127850-01A1) and ARO (W911NF-17-1-0024), to J.K.B. from NIH (GM126904), and to S.H.S., D.K.N and J.K.B. from the Rosen Bioengineering Center at Caltech. E.C.M.T. was supported by a Croucher Foundation Research Fellowship.

## Author contributions

Conceptualization, S.H.S., J.K.B., L.M.T., and D.K.N.; Methodology, S.H.S., E.C.M.T., M.D.Y., F.J.O., S.A.T., E.D.A.S., J.K.B., L.M.T., D.K.N.; Formal Analysis, S.H.S. and L.M.T.; Investigation, S.H.S., E.C.M.T., M.D.Y., F.J.O., S.A.T., E.D.A.S.; Resources, J.K.B., L.M.T. and D.K.N.; Writing – Original Draft, S.H.S. and D.K.N.; Writing – Reviewing & Editing, S.H.S., E.C.M.T., M.D.Y., F.J.O., S.A.T., E.D.A.S., J.K.B., L.M.T., D.K.N.; Visualization, S.H.S.; Supervision, J.K.B., L.M.T. and D.K.N.; Funding Acquisition, J.K.B., L.M.T. and D.K.N.

## Data and materials availability

Data and strains used in this study are available on request. Data and code are available at github.com/DKN-lab/phz_eDNA_2019.

## Materials and Methods

### General

All strains were plated on LB from −80°C glycerol stocks and grown overnight at 37°C. Plates were stored at −4°C for up to a week and were used to inoculate liquid cultures. Liquid cultures were grown in 3mL of medium in glass culture tubes (VWR #47729-583) in an orbital shaker (New Brunswick, Innova 44) at 37°C shaking at 250 rpm.

Chemicals were obtained from commercial sources (Sigma Aldrich, Fisher Scientific, VWR, and New England Biolabs) and used without further purification unless otherwise specified. All solutions were made with Milli-Q water (>18 MΩ cm). Phosphate buffered saline (PBS) used for resuspending cells or contacting biofilms (137mM NaCl, 10mM PO_4_ (1.44 g / L Na_2_HPO_4_, 0.24g / L KH_2_PO_4_), 2.7mM KCl, pH 7.2) was different than PBS used for chemical stocks and in vitro assays (10mM PO_4_ (0.952 g/L Na_2_HPO_4_, 0.592 g/L NaH_2_PO_4_), 50mM NaCl, pH 7.0).

PCA was synthesized by Dr. Stuart Conway’s Lab and was a gift. PCN was obtained from Princeton Biochem (#PBMR030086). PYO was synthesized by shining light on PMS (VWR #AAH56718-06) and purified with repeated organic (dichloromethane) and acid (HCl) extractions as described previously (Costa et al., 2017). PYO was further purified by reverse phase HPLC or repeated hexane precipitations. Stocks of PYO were either made in PBS (pH 7.0) to 1mM or in 20mM HCl to 10mM and quantified by absorbance at 690nm with the extinction coefficient ε690 = 4306 M^−1^ cm^−1^. Stocks of PCA were made in PBS to 1-2mM or in 10mM NaOH to 10mM. Stocks of PCN were made in DMSO to 20mM.

### Colony Biofilms

#### Growth

Colony biofilms were grown on 1% tryptone (BD #211705), 1% agar (BD #214010) in 6-well plates (Fisher Scientific #08-772-49) at room temperature (∼22 °C) in the dark for 3 to 4 days. The medium was autoclaved and cooled to 60°C, then in a biosafety cabinet (The Baker Company #SG603A-HE) 5mL of molten agar was transferred into each well of the 6-well plate and left to solidify for 60min with no lid and constant airflow. When phenazines were added to the medium, concentrated stocks were added first, then the medium, and then the wells were stirred with sterile pipettes until uniform. Membrane filters (Sigma-Aldrich # WHA110606) were gently placed shiny side down onto the center of each agar well, and colonies were inoculated by pipetting 10µL of stationary phase culture (from LB liquid) onto the center of the membranes.

For DNase treated biofilms, after 3 days of growth membranes/biofilms were transferred to fresh agar wells containing 20uL of DNase 1 (2 Kunitz units / µL) (Sigma-Aldrich #D4263) spotted directly underneath the biofilm. Biofilms then grew for another 24 hrs. DNase treatment was done with or without NEB Buffer 4.

#### Imaging

All colony biofilms were imaged in the 6 well plates at 20x zoom with a Keyence digital microscope (VHX-600) before extraction. Colonies were stained for confocal imaging by growing them with 1µM TOTO-1 and 10µM Syto60 in the underlying agar. Biofilms on their membranes were transferred to imaging dishes. High magnification images were taken by gently dropping coverslips (#1.5) onto the top surface of the colony and imaging through the coverslip with an upright confocal microscope, as described below (IDA biofilms – Fluorescence imaging). Colony biofilms on membranes were fixed for scanning electron microscopy (SEM) by floating the membrane on a paraformaldehyde solution and then submerging. Upon submersion, the colony came off of the membrane mostly intact, leaving only a thin layer of cells fixed to the membrane. See SEM imaging section below for further details.

#### Extraction

Membranes, with the colony biofilms stably attached, were lifted off the agar with fine tweezers and carefully placed in microfuge tubes containing 800µL or 1mL of PBS. For colonies that were smaller in diameter than the microfuge tube, the membrane was placed upside down directly on top of the open tube, so that the colony sat above the PBS hanging from the membrane. Then, the colony was gently pushed down into the tube, by pushing with tweezers from the center. If the colony diameter was greater than the tube the membrane/biofilm was carefully held with the tweezers and allowed to flop over. While the membrane/biofilm was curved over it was gently threaded into the tube. If any part of the colony touched the rim of the tube, the tweezers were used to scrape as much as possible into the inside.

Once the membrane/biofilm was in the tube, each tube was vortexed (Fisher Scientific #12-812) at maximum speed for 1 min, after which the vast majority of the biofilm was resuspended in the liquid and no longer associated with the membrane. The membrane was removed from each tube with tweezers. The biofilm suspension was then centrifuged (Eppendorf #5418) at 6000 rcf for 5min. The supernatant was removed and immediately prepared for LC-MS or frozen at −20°C. This process was done for many biofilms at once, so biofilms typically sat for 30-60min in PBS from the time the membrane was first added to the time the samples were centrifuged.

Biofilm volumes were estimated by comparing biofilm suspension volumes to controls that only had bare membranes added and removed. Volumes were measured with a p200 pipette. Significant variation was observed in volume measurements of replicate biofilms that looked identical, so the mean WT colony volume of 60µL was used to normalize all colony measurements, although there are likely subtle volume differences between the differently treated colonies/strains.

The 6-well agar plates were extracted, by adding 2 or 3mL of PBS to the 5mL of agar. The agar was scarred with a pipette tip to facilitate agar/liquid exchange. The 6 well plates were left on an orbital shaker at 250rpm (Ika #KS-260) for 6 hours, which was determined to be optimal. Then, 1mL of the liquid from each well was transferred to a microfuge tube and processed for LC-MS or frozen at −20°C.

For the sonication experiment, resuspended biofilms were treated for 1 min on ice (Fisher Scientific, Sonic Dismembrator 550), then processed normally. CFU counts showed that >65% of cells died. For the no-membrane experiment, 3mL of PBS were added directly to the wells containing the biofilm. The biofilm was quickly resuspended in the liquid using a cell scraper and 1mL of liquid was saved, while the rest was removed. Then the agar was extracted as normal.

#### LC-MS

Samples were filtered with 0.2µm spin filters (Corning #8160) and transferred to sample vials (Waters #600000668CV) and loaded into the autosampler at 10°C. Samples were run on a Waters LC-MS system (Waters e2695 Separations Module, 2998 PDA Detector, QDA Detector). 10µL of each sample was injected onto a reverse phase C-18 column (XBridge #186006035) running a gradient of 100% H_2_O + 0.04% NH_4_OH to 70% acetonitrile + 0.04% NH_4_OH over 11 min (run times were 20min total). UV-Vis and positive MS scans were acquired for each run. PCA, PCN, and PYO were distinguished by retention time (∼3 min, 6 min and 8.85 min respectively), detected at 364 nm (PCA and PCN) or 313 nm (PYO), and manually verified by examining masses 225.2 (PCA), 224.2 (PCN) and 210.24 (PYO). Peaks were automatically identified in the UV-Vis channels by retention time, quantified using the apex track algorithm, converted to concentrations using standard curves (>6 points from 100nM to 100µM), and then exported as text files from the Empower software.

#### eDNA measurements

Resuspended colony biofilms were assayed for eDNA by measuring TOTO-1 fluorescence on a Tecan Spark 10M plate reader in black 96 well plates. Wells were prepared with 65 µL PBS and 10µL of TOTO-1 (10µM stock, 1µM final) and 25µL of biofilm suspension or calf thymus DNA was added to each well and gently mixed by pipetting. TOTO-1 fluorescence was monitored with 480nm excitation and 535nm emission after ∼20 minutes of incubation at room temperature. Colony biofilms were measured with six biological replicates and technical triplicates. A standard curve was obtained by diluting a stock of calf thymus DNA twofold serially, yielding measurements from 500 µg / mL to 7 ng / mL.

### DNA binding assays

Phenazine stocks were made in PBS (pH 7.0 10mM PO_4_, 50mM NaCl) to 1mM (PCA, PYO) or 500µM (PCN with 5% DMSO). All assays were performed in the same PBS buffer unless otherwise noted. Complementary 29bp oligos from IDT were annealed (cooled incrementally over 1 hour following denaturation) to form the ds DNA for ITC and the ethidium displacement assays (see Table S2 for sequences). ds DNA for MST was prepared by PCR amplifying an 80bp region from *P. aeruginosa* genomic DNA using the primers listed in table S2. One of the primers contained a Cy3 fluorophore for the MST readout.

#### Isothermal titration calorimetry (ITC)

ITC was performed with a MicroCal ITC200 (Malvern). Phenazine stocks were loaded directly into the ITC syringe. The cell was loaded with 10 to 50µM ds DNA in PBS (or PBS + 5% DMSO for PCN). Results shown are representative of replicates taken at various ds DNA concentrations. Thermograms were recorded at 21°C with stirring at 300rpm and reference power 2. There were 13 injections of 3.2µL volume spaced by 240 seconds with 600 seconds of settle time.

Thermograms were integrated and baseline corrected in the Origin software. Peak integrations were then loaded into the GUI version of pytc (Duvvuri et al., 2018) with the default 0.1 units of uncertainty added to each measurement. Binding curves were plotted with the molar ratio phenazine / oligomer. The data were fit with the Bayesian model and parameter estimates with confidence intervals were generated. The K value was converted to a dissociation constant by taking the inverse and converted from the oligomer concentration to base-pair concentration by multiplying by 29bp / fraction competent (f_x = 1.6 for PYO and 1.3 for PCN).

#### Ethidium bromide displacement

In black 96 well plates (Nunc #237105) 5µM ethidium bromide (EtBr) was prepared in wells with increasing concentrations of PCA, PCN, or PYO. Fluorescence readings were taken on a Tecan Spark 10M plate reader with excitation 480nm and emission 600nm. Then a small volume of 29bp ds DNA (prepared the same as for ITC) was added to a final concentration of 1µM. The plates were mixed with rotary shaking and incubated at room temperature for at least 5 min protected from light. Then fluorescence was read again, and the bound ethidium signal was calculated by subtracting the pre-DNA reading from the post-DNA reading. IC50 was calculated by fitting the data to the hill equation. IC50 was converted into K_i_ with the equation K_i_ = IC50 / (1 + [EtBr] / K_d_), using an empirical K_d_ for ethidium (under the same conditions) of 1µM.

#### Microscale thermophoresis (MST)

Microscale thermophoresis was performed with a NanoTemper Monolith instrument following the manufacturer’s protocol (capillaries - NanoTemper #MO-K022). Briefly, capillary solutions were prepared with 50nM ds DNA and two-fold dilutions of phenazines starting at concentrations greater than or equal to 1mM. Thermophoresis was performed at an ambient temperature of 22.5°C, with the thermorphoresis laser power 40% and the fluorescence excitation laser at 20% power. Thermophoresis was recorded for 30 seconds and evaluated using the T-jump strategy. Fluorescence peak shapes along the X-axis of each capillary were very uniform and peak intensity did not vary meaningfully between phenazine concentrations. K_d_ was calculated by fitting the quantified data to a hill equation. Results shown are representative of multiple MST runs with varied settings.

#### Endogenous fluorescence of reduced phenazines

The endogenous fluorescence of reduced phenazines was measured with different concentrations of calf thymus DNA with a BioTek Synergy 4 plate reader placed in an anaerobic chamber. Reduced phenazines (100µM solutions) were prepared by bulk electrolysis of oxidized phenazine solutions in electrochemical chambers described previously (Wang et al., 2010). The solutions were transferred into stoppered serum bottles and moved to the anaerobic chamber containing the plate reader. 135µL reduced phenazine was incubated with 15µL PBS containing different amounts of calf thymus DNA, thus each well contained 90µM phenazine. Fluorescence was monitored in a black 96 well plate with filter cubes to control excitation and emission wavelengths. PYO was excited using a 360nm light and emission was monitored at 460nm. For PCA and PCN excitation was at 485nm and emission was at 528nm.

### Phenazine to phenazine electron transfer *in vitro*

Electron transfer reactions between phenazines under anoxic conditions were monitored in the anaerobic plate reader described above. Reduced phenazines (described above) were mixed with oxidized phenazines (67.5µL each) and 15µL PBS with or without DNA (2mg / mL) to yield mixtures containing 45µM of each phenazine. Reactions in clear bottom, black walled 96 well plates were measured at 690nm absorbance approximately one minute following mixing. For each well, an absorbance scan and fluorescence measurements (described above) were also taken, and these data matched the results obtained at 690nm. Time series (2.5 min) of absorbance and fluorescence measurements were taken for each well, but no changes were observed, indicating the reactions had reached equilibrium.

Phenazine redox reactions were monitored over time under oxic conditions with an aerobic Beckman Coulter DU 800 spectrophotometer. The instrument was blanked with PBS. Plastic cuvettes were filled with 500µL PBS (oxic) or PYO_ox_ and placed inside the instrument and the lid was shut.

Reduced phenazine or anoxic PBS was drawn (∼500µL) from stoppered serum bottles (anoxic) into needled 1mL syringes. Absorbance measurements at 690nm (1.5sec intervals) were started on the cuvette and proceeded for 10 seconds to acquire baseline values. Then the lid was opened, and the syringe was quickly emptied into the cuvette and the lid was closed again. The measurement proceeded until 90 seconds had elapsed. Reactions were repeated in triplicate.

### DNA modified electrodes

#### Preparation of thiol-modified oligonucleotide

A single-stranded DNA sequence with the 5ꞌ end modified with a C6 S-S phosphoramidite was purchased from Integrated DNA Technologies (see Table S1 for DNA sequences). The oligonucleotide was reduced using dithiothreitol (DTT, Sigma Aldrich, 100 mM) in a buffer solution (50 mM Tris-HCl, pH 8.4, 50 mM NaCl) for 2 h. The reduced thiol-modified DNA was then purified by size exclusion chromatography (Nap5 Sephadex, G-25, GE Healthcare) with phosphate buffer (pH 7.0, 5 mM NaH_2_PO_4_, 50 mM NaCl) as the eluent. Subsequently, high pressure liquid chromatography (HPLC, HP 1100, Agilent) was performed using a reverse-phase PLRP-S column (Agilent) using a gradient of acetonitrile and 50 mM ammonium acetate (5-15% ammonium acetate over 35 minutes). After HPLC purification, the thiol-modified ssDNA was characterized using matrix-assisted laser desorption ionization (MALDI) characterization using a Autoflex MALDI TOF/TOF (Bruker), and quantified using a 100 Bio UV-visible spectrophotometer (Cary, Agilent). Another ssDNA strand for installing a CC mismatch near the interface between DNA duplexes and self-assembled monolayer (SAM) linker was prepared in an analogous manner as detailed above.

#### Preparation of amine-functionalized oligonucleotide

A single-stranded DNA sequence with the 5ꞌ end modified with a C6 NH_2_ phosphoramidite was purchased from Integrated DNA Technologies. HPLC was performed using a reverse-phase PLRP-S column using a gradient of acetonitrile and 50 mM ammonium acetate (5-15% ammonium acetate over 35 minutes). After HPLC purification, the amine-modified oligonucleotide was characterized using MADLI-TOF-MS and quantified using UV-visible spectrophotometry.

#### Preparation of phenazine-functionalized oligonucleotide

Phenazine-1-carboxylic acid (PCA, 4.9 mg, 22 μmol) was added to dicyclohexylcarbodiimide (DCC, 9.3 mg, 45 μmol) and *N*-hydroxysuccinimide (NHS, 5.2 mg, 45 μmol) in degassed, anhydrous dimethylformamide (DMF, 500 μL) at RT in a scintillation vial wrapped in aluminum foil. The barely soluble PCA turned from green to yellow upon stirring overnight in the dark under Ar. The NHS-activated PCA solution was reduced to low volume (100 μL) and was then added to a solution containing amine-modified DNA (0.42 μmol) in the presence of hydroxybenzotriazole (HOBT, 1 mg, 7.4 μmol) and *N,N,Nꞌ,Nꞌ*-tetramethyl-*O*-(1*H*-benzotriazol-1-yl)uronium hexafluorophosphate (HBTU, 1 mg, 2.6 μmol) in sodium bicarbonate solution (200 μL, pH 8, 0.1 M) overnight in the dark with the 1.5 mL Eppendorf tube wrapped in aluminum foil and mixed thoroughly using the shake function of a benchtop vortexer. The 1.5 mL tube secured using a clip to avoid spilling. The crude product was then buffer exchanged using a NAP-5 size-exclusion column into phosphate buffer (pH 7.0, 5 mM NaH_2_PO_4_, 50 mM NaCl). The treated product was subsequently purified using HPLC on a reverse-phase PLRP-S column with a gradient of acetonitrile and 50 mM ammonium acetate (5-15% ammonium acetate over 35 minutes) while monitoring 252, 260, 280, 354, and 365 nm simultaneously. Pale yellow liquid fractions were collected and freeze-dried on a lyophilizer (Labconco). The resulting yellow dried smear was resuspended in minimal MQ H_2_O and was subsequently analyzed by MALDI-TOF-MS.

#### Formation of double-stranded DNA duplexes

The two strands of a duplex are synthesized separately, purified, desalted, EtOH precipitated, and stored frozen at −20 °C. Prior to electrochemical experiments, the two strands of a duplex were mixed in equimolar (50 μM) in 200 μL phosphate buffer (pH 7.0, 5 mM NaH_2_PO_4_, 50 mM NaCl). The DNA solution was deoxygenated by bubbling Ar for at least 5 minutes per mL, and then annealed on a thermocycler (Beckman Instruments) by initial heating to 90 °C followed by slow cooling over a span of 90 min.

#### Preparation of DNA-modified electrodes

Multiplexed chips are gently cleaned by sonicating with acetone once then isopropanol three times before drying with Ar. They are then cleaned with UV/Ozone using a UVO cleaner for 20 minutes. Immediately after cleaning the surface, a plastic clamp and rubber gasket (Buna-N) were affixed to the Au surface to create a well for liquid and 50 µM duplex DNA in phosphate buffer (5 mM phosphate, pH 7, 50 mM NaCl) to make ds DNA-modified surfaces. The ds DNA was incubated on the surface for 18-24 h in the absence of light. Once the ds DNA is affixed to the surface, it cannot be dried without compromising the structure and subsequently the measured properties of the ds DNA-modified surfaces. The solution was then exchanged 3× with phosphate buffer (pH 7, 5 mM phosphate, 50 mM NaCl, 5% glycerol) and incubated with 1 mM mercaptohexanol in phosphate buffer (pH 7, 5 mM phosphate, 50 mM NaCl, 5% glycerol) for 45 minutes. Lastly the surface was rinsed at least 5× with phosphate buffer (pH 7, 5 mM phosphate, 50 mM NaCl) that was degassed by leaving open in an anaerobic chamber (Coy Lab Products) for at least 3 days.

#### Multiplexed chip electrochemical measurements

Experiments performed were replicated at least three times using different samples, and data presented are from representative trials. Cyclic voltammetry (CV), square wave voltammetry (SWV), and differential pulse voltammetry (DPV) were carried out using a 620D Electrochemical Workstation (CH Instruments) at room temperature inside an anaerobic chamber. The atmosphere of the anaerobic chamber (< 1 ppm O_2_, ca. 3.4% H_2_) was monitored using a CAM-12 O_2_ and H_2_ sensor (Coy Lab Products). The chamber was maintained O_2_-free by using two ventilated Pd catalyst packs (Coy Lab Products).

Electrochemical experiments were carried out in a three-electrode set-up under an anaerobic atmosphere. CV was conducted at a scan rate of 100 mV/s unless otherwise specified. The central well around the multiplexed electrode surfaces created by the plastic clamp was filled with aqueous buffer containing degassed phosphate buffer (pH 7, 5 mM phosphate, 50 mM NaCl). An Ag/AgCl reference electrode (BASi) was coated with a solidified mixture of 1% agarose and NaCl (3 M) in water inside a long, thin pipette tip. The tip was cut so that the salt bridge could connect the electrode to the buffer from the top of the well. A freshly-polished Pt wire used as an auxiliary electrode was also submerged in the buffer from the top of the well. The working electrode contacted a dry part of unmodified gold surface. Scan rate dependence studies (10-5000 mV/s) were carried out to determine whether the phenazine moiety was covalently attached to the DNA-modified electrodes. A linear relationship between scan rates and measured peak currents signified that phenazine was covalently attached. For O_2_ electrocatalysis studies, CV was conducted in open air in O_2_-saturated phosphate buffer (pH 7, 5 mM phosphate, 50 mM NaCl). Hexaammineruthenium(III) chloride (RuHex, 1-500 µM in pH 7, 5 mM phosphate, 50 mM NaCl) was used in control experiments to probe whether electron transfer occurred between the Au electrode and the covalently attached PCN through DNA.

### Time resolved spectroscopy with metal complexes

Colony biofilms (*Δphz\**) were grown and suspensions were prepared in 500µL PBS, as described above for the LC-MS and eDNA measurements. 400µL of the suspension was transferred to a clean quartz cuvette. Time-resolved emission measurements utilized a YAG laser (λexc = 532 nm), with laser powers of ∼2 mJ per pulse. A colored-glass longpass filter (λ > 600 nm) was used to minimize scattered laser light and the emission of the Ru(phen)_2_dppz^2+^ complex was monitored at 620 nm. An excitation pulse (532nm) was delivered and emission was recorded at 620nm for 1.9µs and each timepoint was 0.2ns. A background scan was acquired with only the biofilm suspension. Then 5µM Ru(phen)_2_dppz^2+^ was added and a scan was recorded. Subsequently, 5µM Rh(phi)_2_bpy^3+^ was added after each scan to acquire the 1, 2, 3 and 4 quencher equivalent datasets. This process was repeated for multiple biofilms and a liquid Δphz* culture that was concentrated to the same optical density (500nm) as the biofilm suspensions. Datasets were background subtracted from the biofilm only background scan and were fit with biexponential decay models.

### IDA biofilms

#### Electrode and reactor preparation

Gold IDA electrodes were prepared as previously described (Boyd et al., 2015). IDAs fabricated on glass substrate were ordered from CH Instruments (#012125). A passivation membrane covered the working electrodes except for the overlapping regions that form the 5μm gap and where the leads were attached. Insulated wires (Digi-Key #W7-ND) were attached to the two working electrodes of the IDA with a conductive epoxy (Electron Microscopy Sciences #12670-EE) that was cured at 80°C for 1 hour. A shell for the IDA construct was constructed from 15mL conical vials that were sawed off at the 2.5mL mark. Rough edges were smoothed with a razor blade, and two holes were poked in the bottom of the vial. The IDA with attached wires was placed inside the conical vial and the wires were threaded through the holes at the bottom. A nonconductive epoxy (Amron International #2131-B) was carefully pipetted into the vial with the IDA until it was full to the rim. The epoxy was allowed to set at room temperature in a fume hood for 24-48 hours. IDA constructs were then used immediately or stored in petri dishes and covered to protect from light and dust.

Reactors (Pine Instruments, 265mL water-jacketed electrochemical reactor) used to grow biofilms on the IDA were cleaned thoroughly and autoclaved between uses. For gas control, the ports were stoppered or sealed with o-rings and screw clamps. Gas inflow and outflow lines were attached via 6 inch gassing needles and 16g 1.5in needles, respectively, pierced through separate stoppers. Gas outflow was attached to a liquid overflow vessel, a 500mL Erlenmeyer flask that also had 16g needles as inflow and outflows. The water jacket of each reactor was connected in series to a water chiller (Lytron) to heat the vessels to ∼31°C. Each reactor was filled with 180mL of sterile medium and the IDA construct was suspended from a stopper (size 4) by threading the wires through 16g needles and then removing the needles, so that the electrode was fully submerged in the liquid. IDAs were sterilized in 10% bleach for 30 seconds, then rinsed in sterile medium before submerging in the reactor.

#### Biofilm growth

Biofilms were grown in a minimal medium (MM) with succinate as the carbon source (14.15mM KH_2_PO_4_, 38.85mM K_2_HPO_4_, 42.8mM NaCl, 9.3mM NH_4_Cl, 40mM Na-succinate, adjusted to pH 7.2 and autoclaved then 1x SL-10 trace element solution (Atlas, 2004) and 1mM MgSO_4_).

From a stationary phase MM culture, the reactors with 180mL MM were inoculated to an OD_500_ of 0.005. The medium was exchanged every 24 hours, and biofilms were grown on the IDA for 3 or 4 days in this fashion. During growth, constant temperature was maintained at ∼31°C, reactors were stirred at 250 rpm (VWR #58948-138) and air was bubbled into two reactors at a time with a small aquarium bubbler (Tetra #77851).

Depending on the experiment, biofilms were grown with or without the full electrochemical setup described below. At a minimum, biofilms were grown with the IDA working electrode (disconnected from potentiostat).

#### Electrochemical setup

Electrochemistry was performed using a three electrode setup and CHI 760b or CHI 760e bipotentiostats. The working electrode(s) were on the IDA and connected to wires as described above. The counter electrode was a separate graphite rod (Alfa Aesar #14738) and the reference electrode was a separate Ag/AgCl electrode (BASi #MW-2030 or #MF-2079) assembled as described in the above DNA CT section. Before measurements were made in a reactor, the medium was sparged with N_2_ for at least 10min. Reactors were gently bubbled with N_2_ throughout the measurements. Unless otherwise noted Generator-Collector scans were acquired at 3mV/s, with the collector held at 0 mV. Square wave voltammetry was performed at 300hz with an amplitude of 25mV and an increment of 1mV. Chronoamperometry of metabolic current was acquired at +100mV.

#### Measurement scheme

For experiments where biofilms were soaked in synthetic PYO, biofilms were soaked for at least 10 min. To transfer the IDA biofilm, the potentiostat leads were removed and the reference and counter leads were attached to the transfer reactor electrodes. The IDA biofilm was dipped in fresh medium to remove solution PYO and then submerged in the transfer reactor. The working electrodes were attached, and the open circuit potential was measured to ensure proper connections – this took roughly 30 seconds. For time sensitive experiments, SWVs were immediately recorded. For biofilm D_ap_ and D_loss_ measurements SWV and GC scans were taken consecutively (manually or with a macro) until 15 of each scan had been acquired. For blank D_loss_ measurements, rapid consecutive SWV scans were taken in an automated fashion using the “repeated runs” option in the CHI software. For blank D_ap_ measurements, the blank IDA was incubated with increasing concentrations of PYO in the soak reactor.

#### Fluorescence Imaging

IDA biofilms were prepared for imaging by removing the biofilm coated region of the IDA from the epoxy encased shell. The exposed region of the glass substrate near the epoxy interface was scored with a diamond tipped scribe and then it was snapped off from the rest of the vial/epoxy assembly. With tweezers the IDA fragment was transferred to a slide with a multi-well silicone spacer and dye mixture was added with a pipette directly to the edge of the IDA fragment until it was fully coated. The dye mixture was 10µM Syto60 (Invitrogen #S11342) and 1 or 2µM TOTO-1 (Invitrogen T3600) in PBS. Coverslips (#1.5) were added directly on top of the mixture and sealed with clear nail polish to the spacer.

Slides were then imaged on a confocal microscope. Most imaging was done using an upright Zeiss LSM 880 with Fast Airyscan. Syto60 was excited with a 633nm laser and emission was typically recorded with a bandpass filter from 650 to 750nm. TOTO-1 was excited with a 488 or 514nm laser and emission was typically recorded with a bandpass filter from 530 to 630nm. Tilescans were taken using a 10x objective and high magnification images were taken with a 63x objective. Several images were taken at 63x magnification using the Airyscan module with superresolution settings. For Airyscan images the appropriate filter cubes were used.

Other imaging was done on an inverted confocal Leica model TCS SPE with a 10x objective for tile scanning. Syto60 was excited with a 633nm laser, TOTO-1 was excited with a 488nm laser, and emissions were set by band pass filters as above.

#### Abiotic IDA measurements with Nafion

Electrochemical experiments were recorded with a bi-potentiostat (CHI, Model #760) in a Teflon cell designed in-house to mount gold interdigitated array (IDA) electrodes from CHI. The counter and reference electrodes were a Pt coiled wire and an Ag/AgCl reference electrode. All measurements were recorded at room temperature in 0.2 M Na_2_SO_4_ as the electrolyte purged with Argon. The IDA electrodes were modified with Nafion film by pipetting 20 μL of a 5% wt Nafion solution (Sigma Aldrich) onto the IDA electrodes and allowing a film to form as the solution dried at room temperature. After 2 hours, the modified IDAs were rinsed with ethanol and DI water and then mounted into the base of the Teflon cell.

The Nafion films on the IDAs mounted in the Teflon cell were loaded with various amounts of Ru(NH_3_)_6_ ^3+^ by either exposing the films to a 10 mM solution of Ru(NH_3_)_6_Cl_3_ (Sigma Aldrich) in 0.2M Na_2_SO_4_ at time points from several minutes to several hours or by allowing Ru(NH_3_)_6_ ^3+^ in the films to diffuse out into bulk electrolyte for several hours. The Nafion films were loaded with Ferrocenemethanol (Fc-OH) by exposing the films to a 250 μM solution of Fc-OH in 0.2M Na_2_SO_4_ for several hours. To decrease the amount of Fc-OH in loaded films, the films were allowed to have the Fc-OH in the films to diffuse out into bulk electrolyte for several hours. For each electrochemical measurement after loading or depleting the film, the cell was rinsed with DI water and 2 mL of fresh electrolyte was added. The counter and reference electrodes were position above the IDA electrodes in a Teflon lid, and the electrolyte were purged with Argon for 15 minutes with a blanket of Argon maintained in the head space of the Teflon cell. Three conditions were measured with the Fc-OH - (i) Fc-OH in solution with naked IDAs, (ii) Fc-OH in solution at IDAs modified with Nafion and (iii) Fc-OH loaded into the Nafion modified IDAs.

For the SWV vs. GC method, SWV and GC scans were acquired as described above, and equation 5 (Supp. Text) was used to calculate Dap with the following parameters: A = 0.0213cm^2^, ψ = 0.505, S = 18.4cm, t_p_ = 1/600s. For chronocoulometry, voltage was stepped from 0 mV to −500mV vs Ag/AgCl. From the slope of plots of charge vs. (time)^1/2^, D_ap_ was calculated from the following equation:

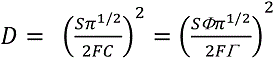

where S is the chronocoulometric slope (C cm^-2^ s^-1/2^), Φ is the film thickness (cm), Γ is the total quantity of Ferrocenemethanol in the film (mol cm^2^), and F is the Faraday constant and C is concentration in moles/cm^3^ ( C = 2.5 x 10^-7^ moles/cm^3^). A film thickness of 10 μm was estimated from the reported density of casted Nafion films by knowing the amount of Nafion deposited and the area covered by the film.

### Scanning electron microscopy (SEM)

IDA and colony biofilms were prepared for SEM using the following protocol. Delicate samples were gently transferred between solutions using metal spatulas or plastic grids as supports. Material was fixed in 4% paraformaldehyde in PBS for 2 hours or overnight, then washed twice with PBS. Material was then fixed in 1% OsO_4_ in water for 1 hour, then washed twice with PBS. Next, the samples were dehydrated by sequentially immersing in 50, 70, 90, 95 and 100% ethanol (EtOH) solutions for 10 min, and then again in 100% EtOH for 1 hour. Samples were then transferred to hexamethyldisilazane (HMDS) solutions of 1:2 then 2:1 HMDS:EtOH for 20min each, followed by two incubations in 100% HMDS for 20min each. Samples were then removed from the solution and air dried before attaching to imaging stubs with conductive tape. Imaging stubs were then sputter coated with 10nm of palladium before being loaded into the SEM. Imaging was done with a Zeiss 1550VP field emission SEM using the SE2 detector.

### Data analysis

Nearly all data processing and analysis was done in R (R Core Team, 2018) using tidyverse packages including ggplot2, dplyr, readr (Wickham, 2017), broom (Robinson and Hayes, 2018) and hms (Müller, 2018). The nonlinear least squares (‘nls’) function in base R was used for D_loss_ fits and the linear model (‘lm’) function was used for D_ap_ fits. Electrochemical data was exported from CHI software as text files with raw data, acquisition parameters and timestamps and then imported into R. Confocal images were stitched (for tilescans) and processed into z-slices or maximum intensity projections using the Zeiss Zen Black software and Fiji (Schindelin et al., 2012).

All code used for data processing and analysis is available in a GitHub repository (github.com/DKN-lab/phz_eDNA_2019). Html versions of the Rmarkdown notebooks are also available as a website (DKN-lab.github.io/phz_eDNA_2019).

