## Supplementary material for "Extracellular DNA promotes efficient extracellular electron transfer by pyocyanin in Pseudomonas aeruginosa biofilms": SI Text and Figures

### Supplementary Text

DNA modified electrode controls

The purpose of carrying out the control experiments shown in Fig. S4 using electrodes modified using self-assembled monolayers (SAMs) of double-stranded DNA (ds DNA) was to probe the electron transfer mechanism between the redox probe and the electrode surface. Four sets of control experiments were conducted to probe whether DNA-mediated charge transport (DNA CT) occurs between PCN and the electrode via the base stack of ds DNA: (i) WM- vs. MM-DNA, (ii) high- vs. low-density monolayers, (iii) PCN vs. RuHex, and (iv) scan rate dependence studies.

(i) WM- vs. MM-DNA: Electrodes modified using SAMs of ds DNA that are well-matched (WM) or contain a single base pair mismatch (MM) were prepared under identical conditions. WM-DNA electrodes and MM-DNA electrodes have similar physical properties but exhibit different charge transfer properties that occur through the aromatic base stacks of DNA. If the electron transfer process between the redox probe and the electrode occurs via DNA CT, then a decrease in the yield of DNA CT is expected due to the perturbation to the base stack introduced by the MM lesion. If the electron transfer process occurs via other modes such as physical diffusion of a charge carrier or physical contact between the redox probe and the electrode surface, then no difference in the current measured would be expected.

(ii) High- vs. low-density monolayers: Electrodes modified using high- and low-density SAMs of ds DNA were used to probe the effect of the number of ds DNA on the charge transport mechanism. A low-density ds DNA monolayer promotes electron transport within one ds DNA, but ds DNA may adopt various conformations because the individual ds DNA are more distantly spaced. A high-density ds DNA monolayer encourages ds DNA to align with each other in a more orderly fashion due to more ds DNA present in a confined space. Regardless of a high- or low-density monolayer, for the case of PCN-modified DNA-SAM, MM discrimination was observed, suggesting that DNA CT is the major mode of electron transport between the electrode and the redox probe.

(iii) PCN vs. RuHex: RuHex is a positively-charged species that binds electrostatically to the negatively-charged phosphate backbone. RuHex also associates to the OH groups at the terminus of the SAM surface passivated by backfilling using mercaptohexanol. Electrons tunnel through the SAM from the electrode to the RuHex situated at the SAM terminus. Electrons can then be transferred and be distributed among the Ruhex bound to the SAM surface and on the DNA phosphate backbone. Residual RuHex present in solution could help with ET through hopping and diffusion in solution, and the contribution of solution-based ET to the current measured depends on the concentration of RuHex used. Therefore a number of RuHex concentrations were screened in order to optimize the conditions for this measurement. For species that do not participate in DNA CT, no MM discrimination will be observed. The RuHex experiments show that the number of ds DNA on the SAM-modified electrodes is similar for both the WM and MM cases. The observation of a sizable MM discrimination demonstrates that PCN bound on DNA participates in DNA CT.

(iv) Scan rate dependence studies (10-5000 mV/s) were carried out to determine whether the phenazine moiety was covalently attached to the DNA-modified electrodes. A linear relationship between scan rates and measured peak currents signified that phenazine was covalently attached.

$D_{ap}$ measurement theory

To measure $D_{ap}$, we compared two electrochemical measurements that depend on $D_{ap}$ in different ways. By performing this comparison at multiple concentrations, we could fit the data points to a line whose slope can be defined by known parameters, yielding $D_{ap}$. *Δphz** biofilms were soaked in 75µM of PYO and then transferred to fresh medium lacking PYO. The biofilm PYO concentration dropped over the course of 45 min as equilibration with the medium occurred. Approximately 15 sets of scans were taken during this time period.

The first measurement was square wave voltammetry (SWV) (Fig. 5B). The SWV peak current ($I_{swv}$) is defined in terms of concentration of redox molecules (*C*) reacting directly (by physical diffusion) or indirectly (through electron self-exchange reactions) with the electrode ($D_{ap}$), the area of the electrode (*A*), the number of electrons per redox reaction (*n*), the Faraday constant (*F*), the pulse width ($t_p$) and a normalization constant ($\varphi$). Note that $I_{swv}$ depends on the square root of $D_{ap}$.

$$I_{swv} = \frac{nFAC\sqrt{D_{ap}}}{\sqrt{\pi t_p}}\varphi \quad (eq. 1)$$

The second measurement was a generator-collector (GC) measurement (Figure 5C). The maximum GC current ($I_{gc}$) is also defined in terms of *C*, $D_{ap}$, *n*, and *F*, but depends on a geometric factor (*S*) (Boyd et al., 2015) as opposed to the electrode area. Note that $I_{gc}$ depends directly on $D_{ap}$.

$$I_{gc} = nFSCD_{ap} \quad (eq. 2)$$

A plot of experimentally determined $I_{gc}$ vs. $I_{swv}$ values yields linear relationships for PYO in biofilms and in solution (i.e., blank IDA) (Fig. 5D, S6), with slope (*m*) that can be defined in terms $I_{gc}$ and $I_{swv}$:

$$m = \frac{I_{gc}}{I_{swv}} = \frac{nFSCD_{ap}}{\frac{nFAC\sqrt{D_{ap}}}{\sqrt{\pi t_p}}\varphi} \quad (eq. 3)$$

$$m = \frac{S\sqrt{\pi t_p D_{ap}}}{A\varphi} \quad (eq. 4)$$

This relationship enables determination of $D_{ap}$ from the experimentally determined dependency of $I_{gc}$ to $I_{swv}$ (i.e., *m*) in terms of known experimental parameters (Fig. 5F). Importantly, it provides a means of determining $D_{ap}$ that is not dependent on knowing C, which for our system is unknown and changes over time as PYO diffuses out of the biofilm.

$$D_{ap} = \frac{(mA\varphi)^2}{S^2 \pi t_p} \quad (eq. 5)$$

Note that because the slope of these plots is linear, it suggests that $D_{ap}$ is constant for a biofilm despite how the concentration of PYO changes as it diffuses. We assume a constant $D_{ap}$ by fitting the data with a linear model.

$D_{loss}$ measurement theory

In order to measure the physical diffusion coefficient of PYO molecules lost from the IDA-grown biofilms, we monitored the loss of biofilm PYO over time as it equilibrated with the fresh medium similar to an approach taken with polymer films (White et al., 1982a). To quantify this process, we used successive SWV scans acquired over 45min following transfer to fresh medium (the SWV subset of the $D_{ap}$ data). A 1-dimensional diffusion model was then applied to fit the decay of $I_{swv}$ yielding an estimate of $D_{loss}$.

Considering 1-dimensional diffusion of an initial mass ($M_0$) of a substance from a point source, the solution of Fick's second law describing the time-dependent concentration gradient is given by eq. 6 where *x* is distance normal from the source, *A* is the cross-sectional area in 3D space, and $D_{loss}$ is the physical diffusion coefficient.

$$C(x,t) = \frac{M_0}{A\sqrt{4\pi D_{loss}t}} e^{-\frac{x^2}{4D_{loss}t}} \quad (eq. 6)$$

When the source is located at no-flux boundary such that the mass diffuses only to one side,

$$C(x=0,t) = \frac{2M_0}{A\sqrt{4\pi D_{loss}t}} \quad (eq. 7)$$

where *C(x = 0, t)* is the time dependent concentration at the surface of the no-flux boundary (e.g. the IDA surface).

To connect this model to the data, the concentration and initial mass can be defined in terms of the measured SWV current ($I_{swv}$). For SWV, the concentration, *C*, is given by:

$$C(t) = \frac{I_{swv}(t)\sqrt{\pi t_p}}{nFA\sqrt{D_{ap}}\varphi} \quad (eq. 8)$$

Concentration can be defined as the mass per volume, so the initial mass, $M_0$, can be expressed in terms of the effective volume probed by the electrode ($V$):

$$M_0 = \frac{VI_0\sqrt{\pi t_p}}{nFA\sqrt{D_{ap}}\varphi} \quad (eq. 9)$$

The initial current is defined as $I_{swv}$ (t = 0) = $I_0$, which is experimentally estimated from the last $I_{swv}$ in the PYO soak, before transfer to PYO-free medium. This is a conservative overestimate, because some of the soak signal comes from the solution PYO. Substituting the values for $C$ (eq. 8) and $M_0$ (eq. 9) into equation 7 yields:

$$I_{swv}(t) = \frac{2I_0V}{A\sqrt{4\pi D_{loss}t}} \quad (eq. 10)$$

The term $V$ refers to the biofilm volume from which the mass of PYO was detected by the electrode. For a 1D electrode process, the concentration gradient extends from the electrode-solution interface ($C = 0$) to the edge of the diffusion layer, $\delta$ ($C = C_{bulk}$) in a near linear fashion (Bard et al., 1980). There is no region where all of the mass is detected, but the electrode has detected one half of the mass in the volume $A \times \delta$, therefore the effective $V$ can be defined:

$$V = \frac{A\delta}{2} \quad (eq. 11)$$

The diffusion layer, $\delta$, for a single potential step can be estimated by:

$$\delta = 2\sqrt{D_{ap}t_s} \quad (eq. 12)$$

where $t_s$ is the amount of time that the driving potential is held. SWV is a series of forward and reverse potential steps for which we could not unequivocally define an equivalent $t_s$ value, however, we discuss reasonable bounds in the assumptions section below. Substituting into equation 11 yields $V = A\sqrt{D_{ap}t_s}$, therefore equation 10 can be written as:

$$I_{swv}(t) = \frac{I_0\sqrt{D_{ap}t_s}}{\sqrt{\pi D_{loss}t}} \quad (eq. 13)$$

And with that expression we can fit the decay of $I_{swv}(t)$ to a model of the form:

$$y = \frac{b}{\sqrt{t}} + a \quad (eq. 14)$$

where the coefficient, *b*, is described in known variables except for $D_{loss}$ and where *a* accounts for any constant background signal.

$$b = \frac{I_0\sqrt{D_{ap}t_s}}{\sqrt{\pi D_{loss}}} \quad (eq. 15)$$

Therefore, $D_{loss}$ can be calculated from the fit data as (Fig. 5F, S7):

$$D_{loss} = \frac{{I_0}^2 D_{ap} t_s}{\pi b^2} \quad (eq. 16)$$

*Model assumptions*

The $D_{loss}$ analysis described above is based on a number of assumptions. It assumes, for example, that the biofilms are homogeneous. In reality, they are heterogeneous, containing many voids and obstacles (e.g., cells and exopolysaccharides) through which diffusion would not occur. We contend heterogeneity would affect $D_{loss}$ and $D_{ap}$ in a similar manner, as it reduces the biofilm volume in which PYO resides. As such, we don't expect biofilm heterogeneity to greatly affect the determination of $D_{loss}$ from $D_{ap}$. Importantly, our analysis assumes a single infinitely thick phase described by a single $D_{loss}$. In reality, there are at least two phases, a thin biofilm adjoining an infinitely thick solution. If $D_{loss}$ in solution is greater than in the biofilm, then at any instance the concentration gradient of PYO across the biofilm will be steeper than predicted by the model. Since the flux of PYO out of the biofilm at any instance is proportional to the product of the gradient and $D_{loss}$, as the model fits the rate of change of PYO in the biofilm the assumption of a shallower gradient than the actual gradient is expected to result in a calculated $D_{loss}$ that is higher than the actual $D_{loss}$.

To estimate the scan time parameter, $t_s$, we assumed that for the blank IDA $D_{ap} = D_{loss}$ and solved for $t_s$ that best fit the $I_{swv}$ decay for the blank IDA. We then used this value, $t_s$ = 21ms, to calculate $D_{loss}$ for the biofilm IDAs. The scan time parameter is intended to estimate the thickness of the diffusion layer that is formed during a single potential step that drives the flux of the electrode reactant resulting in the observed current. SWV is, however, a series of forward and reverse potential steps superimposed on a series of forward potential steps. Therefore, determining the effective scan time that describes the change in thickness of the diffusion layer that occurs during the forward pulse contributing to $I_{swv}$ is nontrivial (we only consider the forward pulse as the reverse scan partially replenishes the electrode reactant that is depleted during the forward scan). One approach to estimate $t_s$ is setting the SWV expression equal to twice the Cottrell equation (eqs. 17 & 18), since $I_{swv}$ is the net current from the forward and reverse pulses. For a pulse amplitude of 0.025 V and step increment of 0.001 V as used here, each potential step is effectively 0.05 V. The potential at which the $I_{swv}$ occurs is the formal potential of the electrode reactant and applying the Nernst equation, the fraction of electrode reactant in the oxidized state at the electrode surface at the start of the forward potential step generating $I_{swv}$ (E = Eo'+ 0.025 V) is 82.6% and at the end (E = Eo -0.025 V) is 17.4%. The Cottrell equation assumes the potential step drives a redox reaction in which 100% of the electrode accessible redox molecules at the electrode go from oxidized to reduced (or vice versa). Replacing C in the Cottrell equation with 0.655 x C to reflect the fraction of PYO at the electrode surface that changes oxidation state during the forward potential step of the SWV yields an estimate $t_s \approx$ 6ms.

$$I_{swv} = \frac{nFAC\sqrt{D_{ap}}}{\sqrt{\pi t_p}}\varphi = 2I_{ps} = \frac{nFA\,0.655 \times C\sqrt{D_{ap}}}{\sqrt{\pi t_s}} \qquad (eq. 17)$$

$$t_s = \frac{t_p\,(2 \times 0.655)^2}{\varphi^2} \qquad (eq. 18)$$

As such, our $t_s$ estimate (21 ms) based on the blank $D_{ap} = D_{loss}$ used to estimate $D_{loss}$ for the biofilm may be an overestimate. Noting that $D_{loss}$ scales linearly with $t_s$ (eq. 16), this would conservatively underestimate the difference between biofilm $D_{ap}$ and $D_{loss}$.

Parameters for electrochemical calculations

The surface area of the electrode for SWV, $A$ = 0.025cm$^2$, and the geometric factor for GC, $S$ = 18.4cm, were calculated for a blank IDA using the known concentration and $D_{ap}$ for ferrocene methanol (Boyd et al., 2015). All quantified SWVs were acquired with a pulse amplitude of 25mV at a frequency of 300Hz and an increment of 1mV. The SWV pulse time, $t_p$, is one half the square wave period (½ * 1/300 = 1/600 sec). Peak separation from CV of PYO in solution indicated that it did not undergo the full 2 electron reduction, but on average underwent electron transfer with $n \approx 1.8$. From these acquisition parameters, $\varphi = 0.7$ was inferred from a table of existing values (Lovrić, 2010). For the $D_{loss}$ estimate, it was assumed that $I_0$ was the soak SWV peak current. The equivalent scan time, $t_s$, for the $D_{loss}$ calculation is discussed above.

### Supplemental Figures

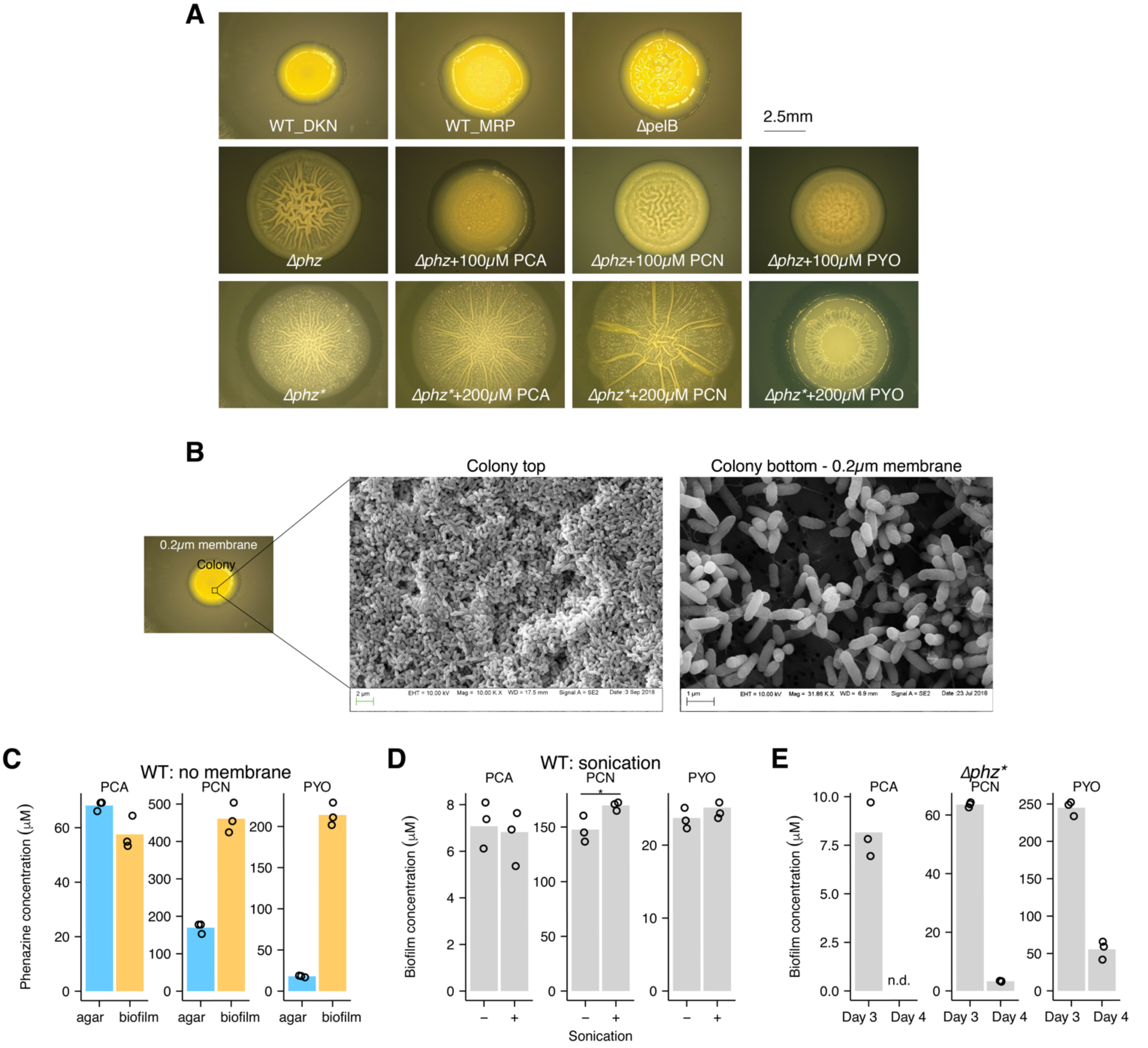


**Figure S1.** Colony biofilms images and controls. (A) Representative images of colony biofilms formed by WT-DKN, WT-MRP, Δ*pelB*, and Δ*phz* and Δ*phz** grown with each phenazine. (B) SEM image of cells at the top and bottom (attached to the 0.2μm membrane) of the colony biofilm. (C) LC-MS quantification of phenazines from colony biofilms grown without an underlying membrane. (D) Comparison of phenazines from WT colonies that were lysed with sonication or not. Statistical test was a Welch's single tailed t-test and the star denotes $p<0.05$. (E) Accumulated phenazine from three *Δphz** colony biofilms following three days of growth with synthetic phenazine (Day 3), and one day later after transfer to fresh agar (Day 4). PCA was not detected on Day 4. Same data are shown in Fig. 1 H.

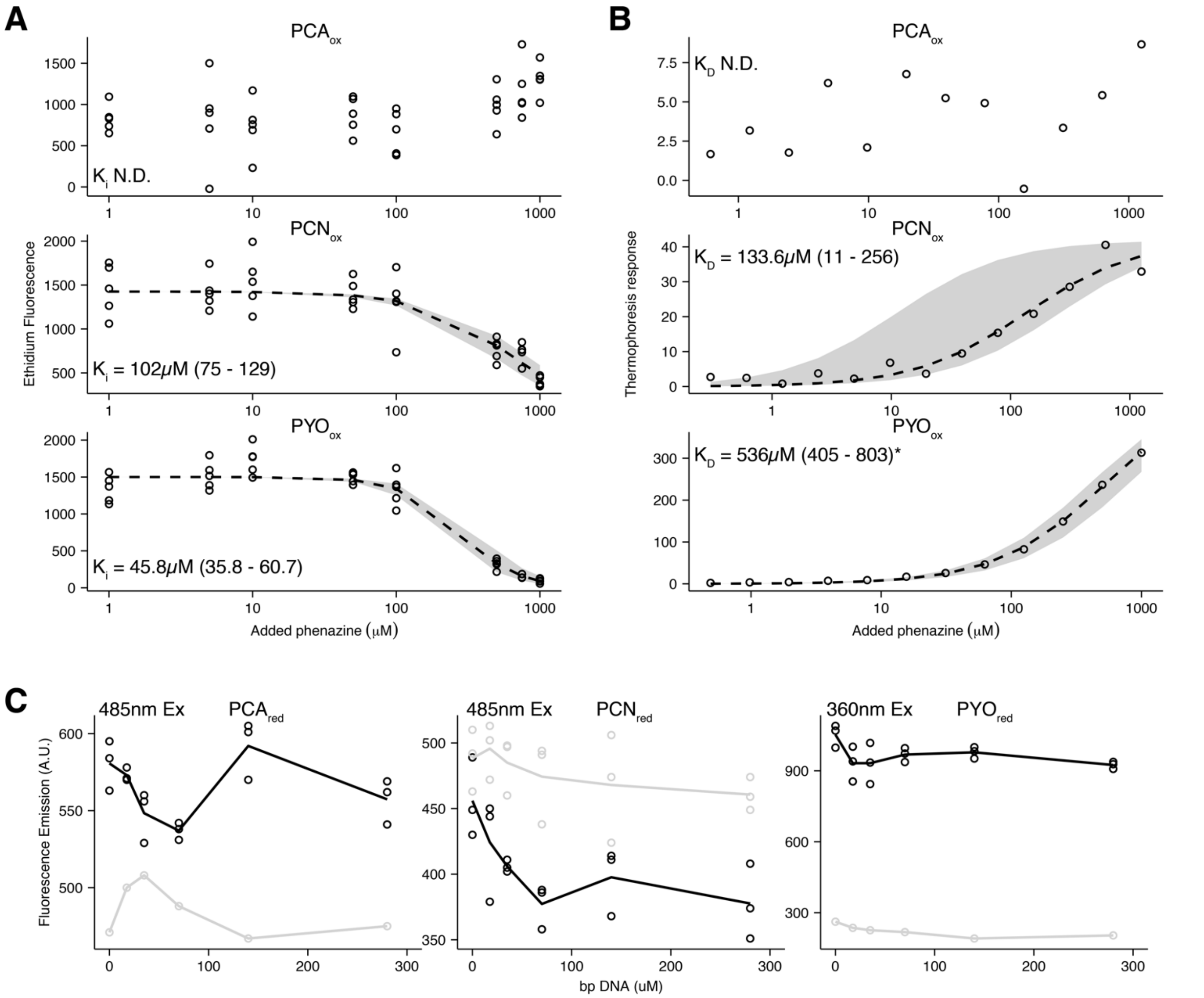


**Figure S2.** Phenazine – DNA binding assays. (A-B) Dotted lines show best fit Hill equations with shaded regions showing curves from the 95% confidence intervals for IC50 or $K_D$. (A) Ethidium bromide displacement from 1uM 30bp ds DNA by oxidized phenazines, as measured by change in ethidium fluorescence before and after incubation with DNA. Assays were done with 5uM ethidium, which has a $K_D$ of 1uM under these conditions. $K_i$ for phenazines was calculated from the relationship $K_i$ = IC50 / (1 + [EtBr] / $K_D$). (B) Microscale thermophoresis binding assay of three oxidized phenazine derivatives with 50nM 80bp cy3 tagged ds DNA. *PYO elicited a strong thermophoresis response that did not saturate, therefore the calculated $K_D$ is likely not relevant. (C) Endogenous fluorescence of reduced phenazines at increasing DNA concentrations. Black lines show experimental conditions and gray lines show buffer only control wells. $PCN_{red}$ did not show fluorescence above background, but the values were reported to show that adding DNA did not facilitate fluorescence emission.

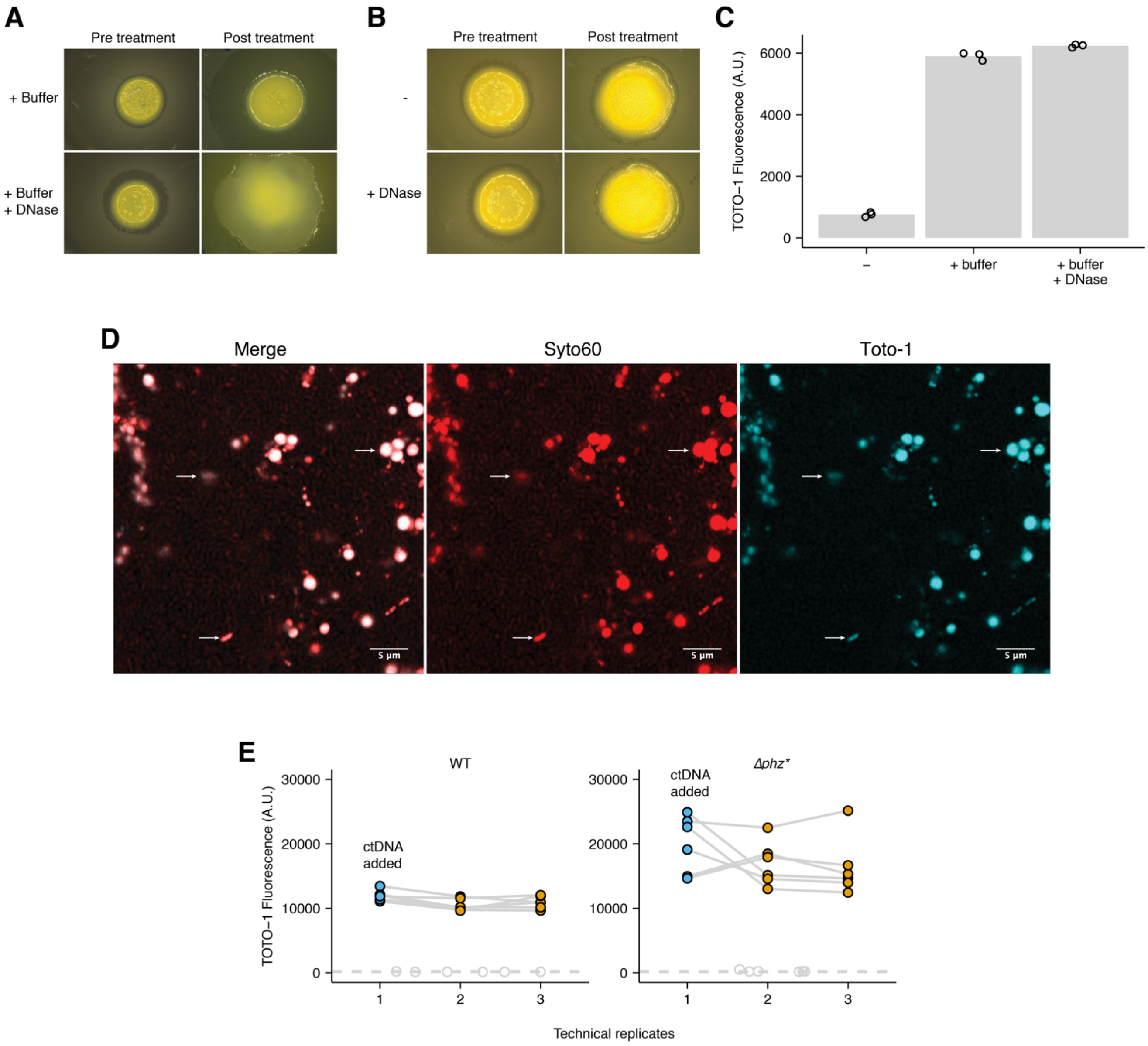


**Figure S3.** eDNA in colony biofilms. (A) Images show representative WT colony biofilms on Day 3 (pre-treatment) and Day 4 (post treatment) with DNase and buffer or only buffer (NEB Buffer 4). (B) DNase treatment of colonies without buffer corresponding to phenazine measurements shown in Fig. 2B. (C) Quantification of eDNA (i.e. cell death) in colony biofilm extracts from A measured by TOTO-1 fluorescence in a plate reader. (D) A high magnification confocal image showing different eDNA structures at the surface of a colony biofilm (white arrows). (E) TOTO-1 measurements of WT and Δ*phz** colony biofilm suspensions. Blue dots show technical replicates where 160ng of calf thymus DNA were added to assess the sensitivity of the assay. Dotted lines and gray dots show background fluorescence values of technical replicates acquired without the dye.

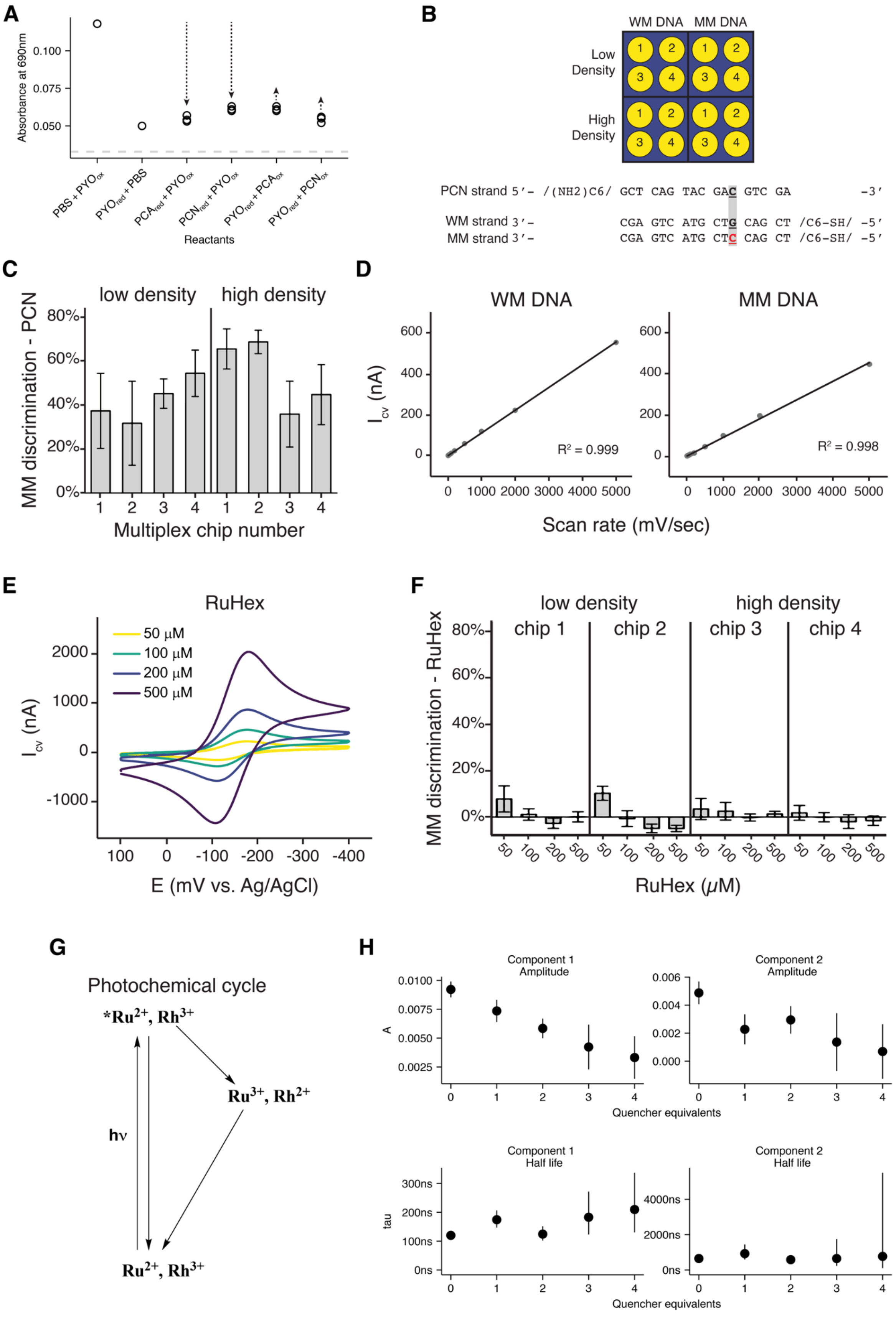
A
Absorbance at 690nm
0.100
0.075
0.050
Reactants
B
WM DNA
MM DNA
Low Density
High Density
PCN strand 5'– /(NH2)C6/ GCT CAG TAC GAC GTC GA -3'
WM strand 3'– CGA GTC ATG CTG CAG CT /C6-SH/ -5'
MM strand 3'– CGA GTC ATG CTC CAG CT /C6-SH/ -5'
C
low density
high density
MM discrimination - PCN
Multiplex chip number
D
WM DNA
MM DNA
R² = 0.999
R² = 0.998
Scan rate (mV/sec)
E
RuHex
50 µM
100 µM
200 µM
500 µM
E (mV vs. Ag/AgCl)
F
low density
high density
chip 1
chip 2
chip 3
chip 4
MM discrimination - RuHex
RuHex (µM)
G
Photochemical cycle
*Ru2+, Rh3+
Ru3+, Rh2+
Ru2+, Rh3+
hν
H
Component 1 Amplitude
Component 2 Amplitude
Quencher equivalents
Component 1 Half life
Component 2 Half life
tau
Quencher equivalents

**Figure S4.** DNA modified electrode details. (A) Inter-phenazine electron transfer reactions in solution, including PYO as the reductant for $PCA_{ox}$ and $PCN_{ox}$. Dashed line shows the background signal from PBS alone or with PCN or PCA. (B) Top - layout of the multiplex chip electrodes used for the measurements. Bottom - the oligomer sequences used to assemble the PCN and thiol modified ds DNA monolayers. (C) Mismatch (MM) discrimination for the high density and low-density DNA monolayers from four multiplex chips of the layout shown in B. MM discrimination was calculated by comparing peak integrations from WM and MM electrodes using the formula 1 - (avg. MM / avg. WM) × 100%. Error bars are standard error propagated from the four WM and four MM electrodes. (D) Scan rate dependence of both the well matched and mismatched surfaces showed linear dependence with increasing scan rate, consistent with a bound redox species. (E) Cyclic voltammetry of hexaammineruthenium(III) chloride (RuHex), which does not participate in DNA CT. Signal is proportional to DNA surface concentration. (F) MM discrimination for different RuHex concentrations calculated in the same way as C. (G) The detailed photochemical cycle referred to in Fig. 3G. (H) The biexponential fit coefficients from Fig. 3I. Error bars show two standard deviations.

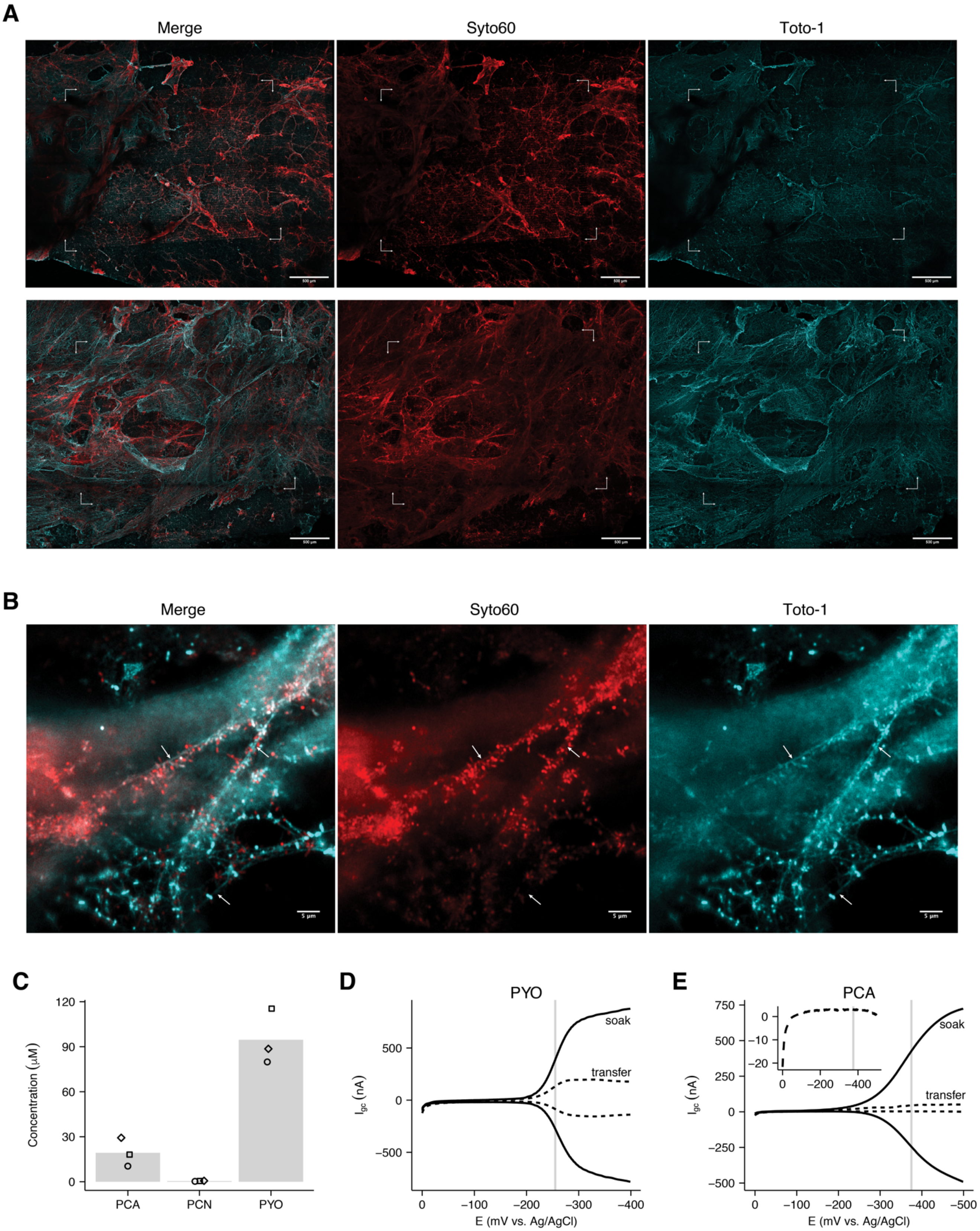
A
Merge
Syto60
Toto-1
B
Merge
Syto60
Toto-1
C
Concentration (μM)
120
90
60
30
0
PCA
PCN
PYO
D
PYO
soak
transfer
$I_{gc}$ (nA)
500
0
−500
0
−100
−200
−300
−400
E (mV vs. Ag/AgCl)
E
PCA
750
500
250
0
−250
−500
0
−10
−20
0
−200
−400
soak
transfer
$I_{gc}$ (nA)
0
−100
−200
−300
−400
−500
E (mV vs. Ag/AgCl)

**Figure S5.** IDA biofilm characterization. (A) Confocal microscopy tilescan images of the two IDA Δ*phz** biofilms used for $D_{ap}$ and $D_{loss}$ analysis. Brackets show the bounds of the IDA working electrode array. Images were acquired at 10x magnification as a tilescan zstack. The stitched maximum intensity projections are shown. (B) Images show individual cells and fine eDNA structures (arrows) at high magnification (63x with digital zoom and airyscan processing). Images are from a single slice of a zstack. (C) LC-MS quantification of WT culture supernatant from an IDA reactor on three consecutive days before medium exchange. (D) Generator collector measurements of Δ*phz** biofilms soaked with PYO. Measurements were acquired during the soak and immediately following transfer to a reactor with fresh medium. (E) Same measurements as D, except the Δ*phz** biofilm was soaked in PCA. The inset figure shows as the collector current with more detail.

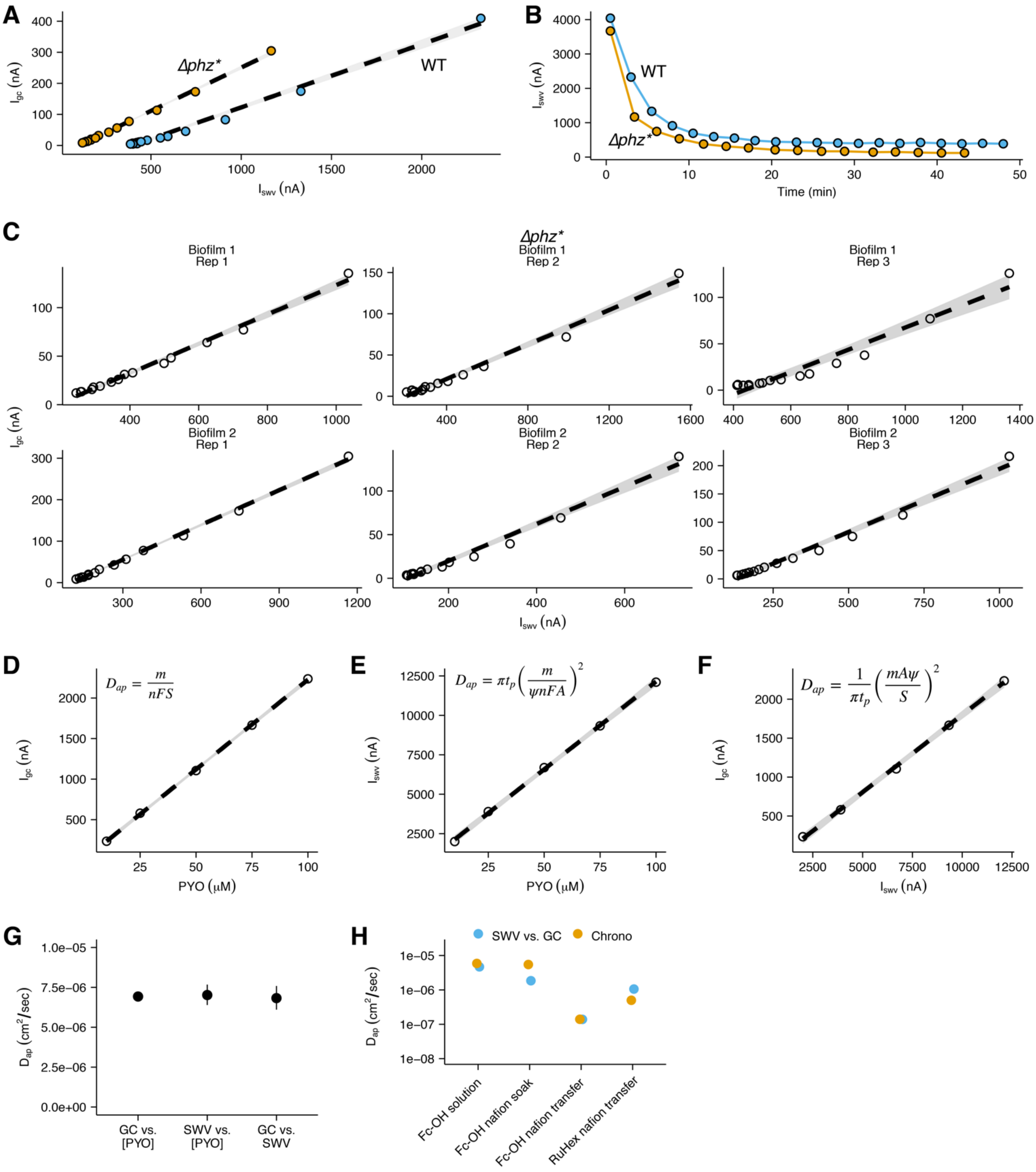
A
B
C
D
E
F
G
H
Δphz*
WT
$I_{gc}$ (nA)
$I_{swv}$ (nA)
Time (min)
Biofilm 1
Rep 1
Rep 2
Rep 3
Biofilm 2
$D_{ap} = \frac{m}{nFS}$
$D_{ap} = \pi t_p \left( \frac{m}{\psi nFA} \right)^2$
$D_{ap} = \frac{1}{\pi t_p} \left( \frac{mA\psi}{S} \right)^2$
PYO (μM)
$D_{ap}$ ($cm^2$/sec)
GC vs. [PYO]
SWV vs. [PYO]
GC vs. SWV
SWV vs. GC
Chrono
Fc-OH solution
Fc-OH nafion soak
Fc-OH nafion transfer
RuHex nafion transfer

**Figure S6.** IDA $D_{ap}$ measurements. (A) Comparison of WT and Δ*phz** + PYO IDA measurements used to determine $D_{ap}$. (B) Comparison of WT and Δ*phz** retention of PYO signal over time. (C) Linear fits for $D_{ap}$ analysis for two Δ*phz** IDA biofilms. Dashed lines show best fit linear models for each subset of data. Shaded regions show 95% confidence intervals from the linear models. (D) GC peak current vs. PYO concentration. The slope, m, is used to define $D_{ap}$ as shown in the expression. The dashed line is a linear fit through the data points and the gray region is a 95% confidence interval. (E) SWV peak current vs. PYO concentration in the same format as D. (F) GC peak current vs. SWV peak current in the same format as D. (G) Estimates of $D_{ap}$ from the three methods shown in D-F with 95% confidence intervals. See supplemental text for the parameter values used. (H) $D_{ap}$ estimates from abiotic IDA experiments with or without the polymer, Nafion, and RuHex or Fc-OH redox molecules.

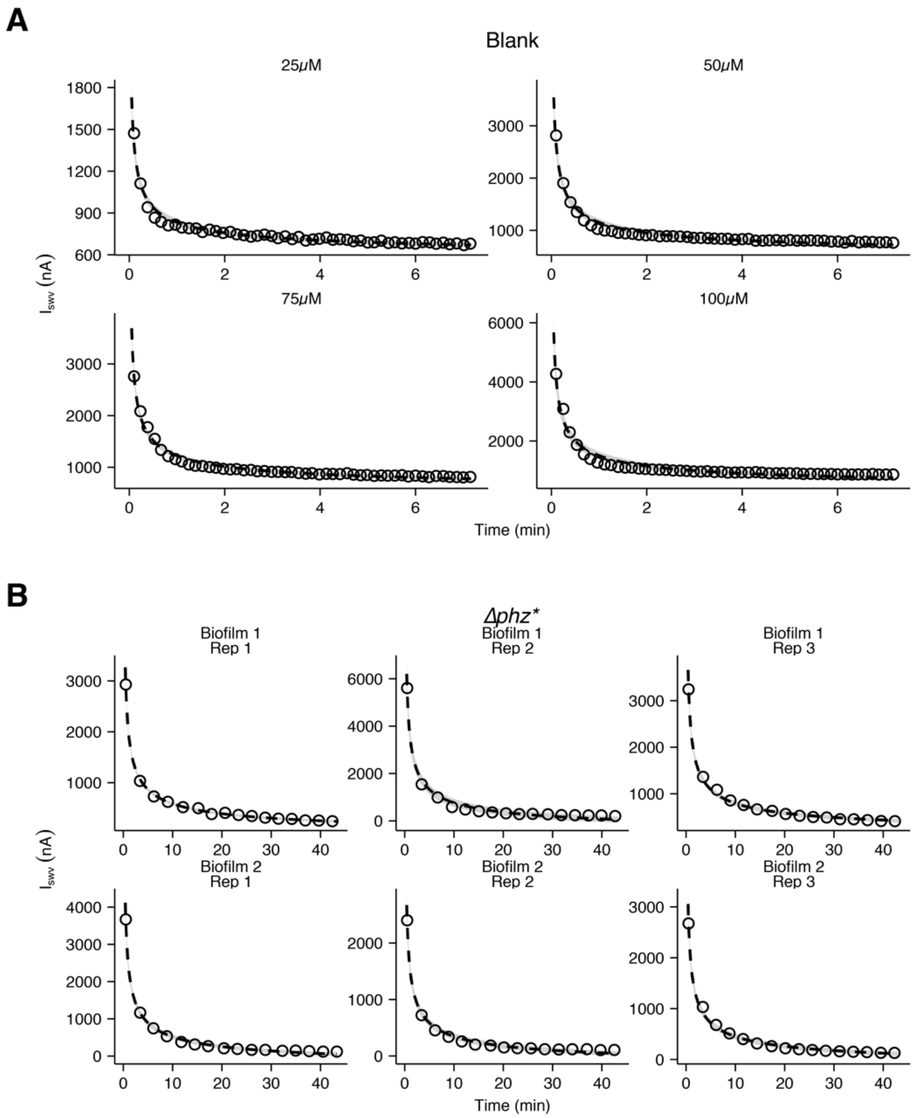


**Figure S7.** IDA $D_{loss}$ measurements. (A) Nonlinear fits for $D_{loss}$ analysis of the blank IDA with different concentrations of PYO. Dashed lines are best fit models. Shaded regions show curves generated from the 95% confidence intervals for the parameter estimates. (B) Nonlinear fits for $D_{loss}$ analysis of two Δ*phz** biofilms (three technical replicates each). Dashed lines are best fit models. Lines and shaded regions same as above.

| Strain name | Genotype | Description |
|---|---|---|
| WT_DKN | | *Pa* PA14 WT from Newman Lab. |
| Δ*phz* | Δ*phzA1-G1* Δ*phzA2-G2* | Mutant incapable of synthesizing PCA. Derivative of WT_DKN. |
| Δ*phz** | Δ*phzA1-G1* Δ*phzA2-G2* Δ*phzMS* Δ*phzH* | Mutant incapable of synthesizing PCA or modifying exogenous PCA. Derivative of WT_DKN, from Dietrich lab. |
| WT_MRP | | *Pa* PA14 WT from Parsek Lab. |
| Δ*pel* | Δ*pelB* | Mutant incapable of synthesizing Pel polysaccharide. Derivative of WT_MRP. |

Strains are all derivatives of *Pseudomonas aeruginosa* UCBPP-PA14 (Schroth et al., 2018).

**Table S1.** Strains used in this study.

| Assay | Description | Sequence (5’ – 3’) |
| --- | --- | --- |
| ITC 29bp duplex | Top | GTGGCAGGTCAGTCAAGTATACTGCACTA |
| | Bottom | TAGTGCAGTATACTTGACTGACCTGCCAC |
| MST PCR 80bp fragment | Forward primer | Cy3/AGAGCAGTTGGTCAAGGC |
| | Reverse primer | GAAAATAACGCTTGACGGAA |
| DNA mod electrode duplex | Top - PCN | $(NH_2)$-$C_6$/GCTCAGTACGACGTCGA |
| | Bottom WM | (SH)-$C_6$/TCGACGTCGTACTGAGC |
| | Bottom MM | (SH)-$C_6$/TCGACCTCGTACTGAGC |

**Table S2.** DNA sequences used in this study.
